# Multiple full-length homozygous IGH haplotypes from Mauritian cynomolgus macaques

**DOI:** 10.1101/2024.11.27.625687

**Authors:** Simone Olubo, William S. Gibson, Trent M. Prall, Julie A. Karl, Roger W. Wiseman, David H. O’Connor, Daniel C. Douek, Chaim A. Schramm

## Abstract

**Background:** Nonhuman primates are valuable experimental models for human disease pathology and vaccine design. However, the vast and mostly uncatalogued immunogenomic diversity of typical species adds complexity to the interpretation of experiments and hinders reproducibility. Mauritian cynomolgus macaques (MCM) offer a unique opportunity to circumvent these difficulties, due to their restricted genetic diversity.

**Results:** We assembled high-quality immunoglobulin heavy chain (IGH) haplotypes from long-read genomic sequencing of 13 MCM. Four animals were homozygous for IGH, yielding 3 distinct haplotypes, termed H1, H2, and H3. IGH haplotype H1 was observed in two of the homozygotes and 5 additional heterozygous animals, accounting for half of the assemblies recovered. H1 shares only 83% average sequence identity with the IGH locus of the rhesus macaque reference genome, in addition to numerous large structural variations. The other two homozygous haplotypes exhibited considerable variation, including a 60 kilobase (Kbp) deletion and 200 Kbp insertion relative to H1. Furthermore, we annotated the IG gene content from all complete MCM IGH assemblies and found 288 functional IGHV alleles, of which 94 (33%) were not in existing databases. We also identified 68 functional IGHD alleles, 11 functional IGHJ alleles, and 33 functional constant gene alleles across all 5 isotypes.

**Conclusions:** We identified multiple common and genetically diverse IGH haplotypes within MCM and provide high-quality reference assemblies and annotations for these to facilitate future work with this important animal model.

## Background

Cynomolgus (*Macaca fascicularis*) and rhesus (*Macaca mulatta*) macaques share a close evolutionary history with humans and are widely used as biomedical models of human health [1–7]. Mauritian-origin cynomolgus macaques (MCMs) descend from a small (∼20) founding population introduced to the island of Mauritius in the 16th century. As a result of this bottleneck event, MCMs exhibit significantly constrained genetic diversity compared to other cynomolgus and rhesus macaques [8]. This makes today’s population of more than 50,000 MCM [8,9] excellent model organisms especially in the context of immunological studies where germline variation can significantly impact results [10–12].

Immune loci such as the major histocompatibility complex (MHC) [13–18], killer cell immunoglobulin-like receptors (KIR) [19,20], and Fc-gamma receptors (FcγR) [21] all have limited genetic diversity in MCM. Relatively little is known about the T cell receptor (TR) [11,22–24] and IG loci [11,25,26] in MCM. The high density of segmental duplications and repetitive elements, as well as the presence of large structural variations, make accurate assembly of these loci difficult [25,27,28]. The IGH locus is particularly challenging and requires specialized sequencing and assembly methods alongside purpose-built annotation tools [27–33]. B cells produce receptors (BCR) and antibodies (Ab) to neutralize or destroy viruses, bacteria, aberrant cells, and other antigenic material. The generation of a diverse Ab repertoire relies on somatic rearrangement of genomic DNA in an individual B cell to create a complete BCR/Ab sequence, a process called V(D)J rearrangement. In the germline, individual IG genes, including variable (V), diversity (D), and joining (J) genes, are selected and joined together permanently altering the resident B cell’s genome. V(D)J rearrangement in the IG heavy chain locus (IGH) generate a VDJ sequence corresponding to the variable region of an Ab heavy chain. The light chain loci, kappa and lambda, do not contain D genes and generate VJ recombination sequences which encode the Ab light chain variable region [34]. The genetic content and organization of the IG loci impact which genes are used most frequently in functional repertoires, and thus are of great interest for immunological studies [35–39].

As an alternative to genomic sequencing, the germline sequences of variable (V), diversity (D), and joining (J) genes can be inferred from adaptive immune receptor repertoire sequencing (AIRR-seq) [40–43]. The Karolinska Macaque database (KIMDB) [26,27] contains 615 and 14 functional IGHV and IGHJ alleles, respectively, inferred from 18 cynomolgus macaques of Indonesian and Mauritian origin. Of these, 279 IGHV and 11 IGHJ alleles were found specifically in MCM. Another recent study added an additional 84 inferred IGHV alleles from 3 Chinese-origin cynomolgus macaques [11]. The large number of reported IGH alleles from a relatively small number of sequenced individuals suggests the macaque IGH loci harbor significant diversity, most of which has not yet been characterized as is true for many other species [35,44,45].

Despite these successes, AIRR-seq-derived inferences have limited ability to resolve V, D, and J genes due to the removal of germline nucleotides during V(D)J rearrangement from the 3’ end of V genes, 3’ and 5’ end of D genes, and 5’ end of J genes. Additionally, most AIRR-seq-derived sequences capture only a small fraction of the constant (C) gene and are unable to reveal C gene polymorphisms which can impact function [46,47]. Furthermore, large structural variations within the IGH locus have been observed in macaques [25,27,29], similar to other primates [28], and humans [35]. High quality genomic references are thus essential for elucidating the complexity of the IGH loci in MCMs and accurate annotation of rearranged IGs. Additionally, the power of the MCM as a model organism relies on a comprehensive understanding of genetic diversity within the constrained population.

While there are multiple high quality, annotated assemblies of the IG loci for rhesus macaques [25,27,29,48], the only genomic IG reference for cynomolgus macaques is an unlocalized scaffold from a 2013 draft genome primarily generated using 2x100 bp paired-end short reads [49]. A new telomere-to-telomere cynomolgus reference genome has recently become available [50] but IG genes have not been annotated. In addition, there are known to be differences in IGH germline alleles and gene usage between MCM and other cynomolgus populations [26]. To fill in this gap, we leveraged long-read whole-genome sequencing data from 13 MCMs [20] to generate curated and annotated assemblies of the MCM IGH locus. Despite the genetic restriction of MCMs, we identified significant allelic and structural variation between haplotypes further demonstrating the potential of MCMs as a powerful model for immunogenetic investigation and reproducibility.

## Results

### Efficient assembly of IGH from whole genome long read sequencing

Long read technologies enable the assembly of highly repetitive regions and the resolution of structural variants, including IGH [28,30,51]. We previously generated whole genome sequencing of 13 MCMs using a combination of Oxford Nanopore Technology and PacBio HiFi sequencing [20]. We assembled the reads for each animal using Hifiasm [52,53] and extracted the IGH locus from contigs that either partially or completely spanned the region. We recovered 13 scaffolds from 10 animals that spanned the entire IGH locus from IGHA to the most distal IGHV7 gene (Table 1). Notably, four animals were homozygous in IGH (Fig 1a and Fig S1).

**Table 1.**
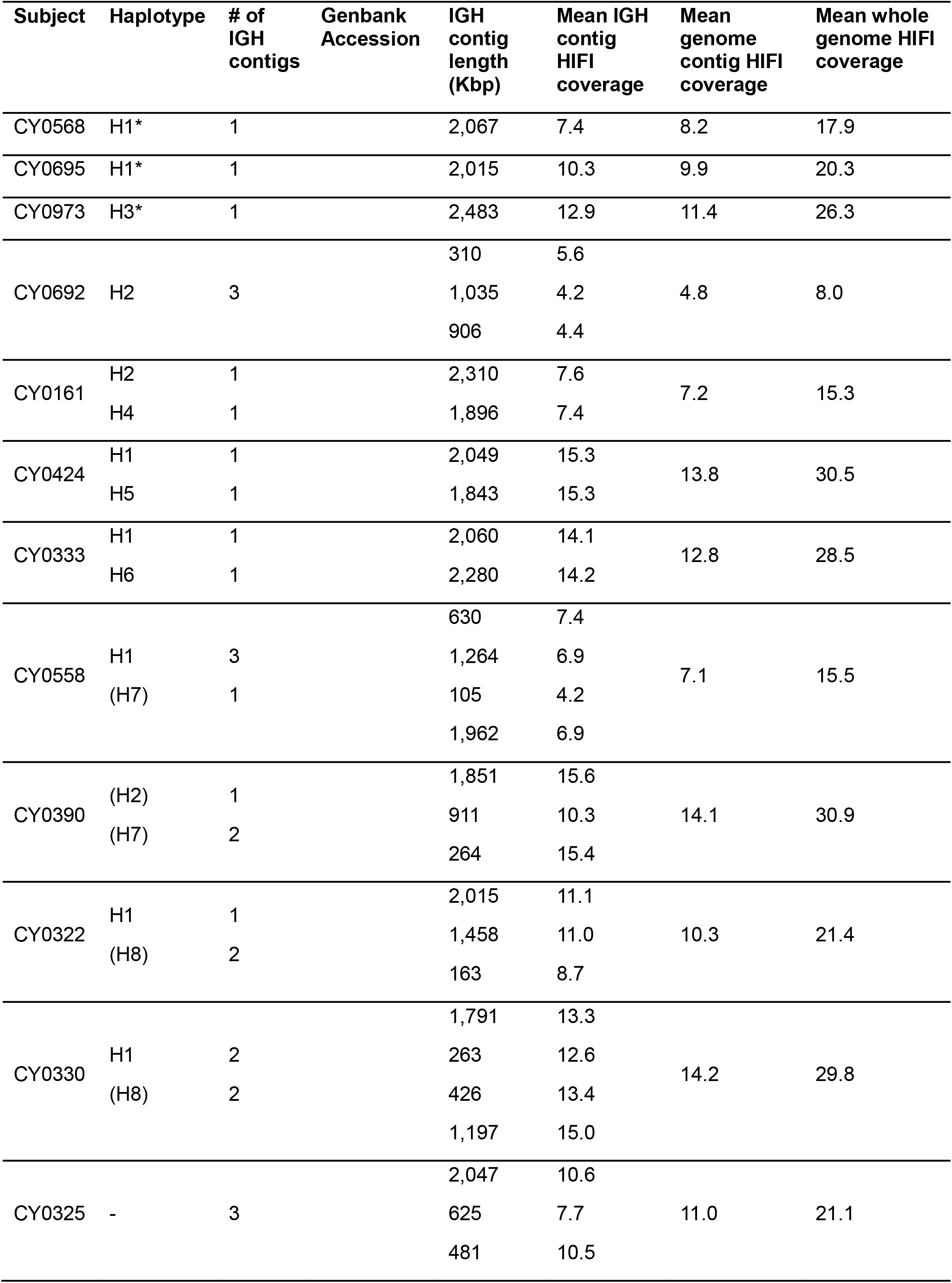
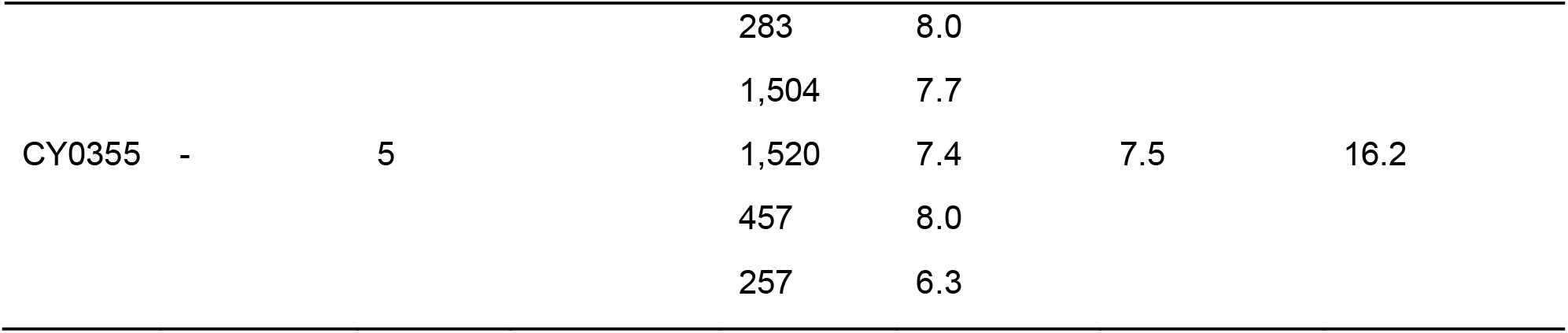
IGH contigs assembled from long-read whole genome sequencing of 13 Mauritian-origin cynomolgus macaques. Asterisk denotes homozygous contig that completely spans IGH. Partial assemblies for a single haplotype are ordered from centromeric to telomeric. Parentheses indicate unscaffolded partial assemblies. Dash denotes unclassified or unresolved haplotype. IGH contig coverage is the mean read depth for each contig, contig coverage is the mean read depth across all contigs, and whole genome coverage is calculated as total base pairs mapped divided by 3 Gbp (the approximate size of the MCM genome).

**Figure 1.**
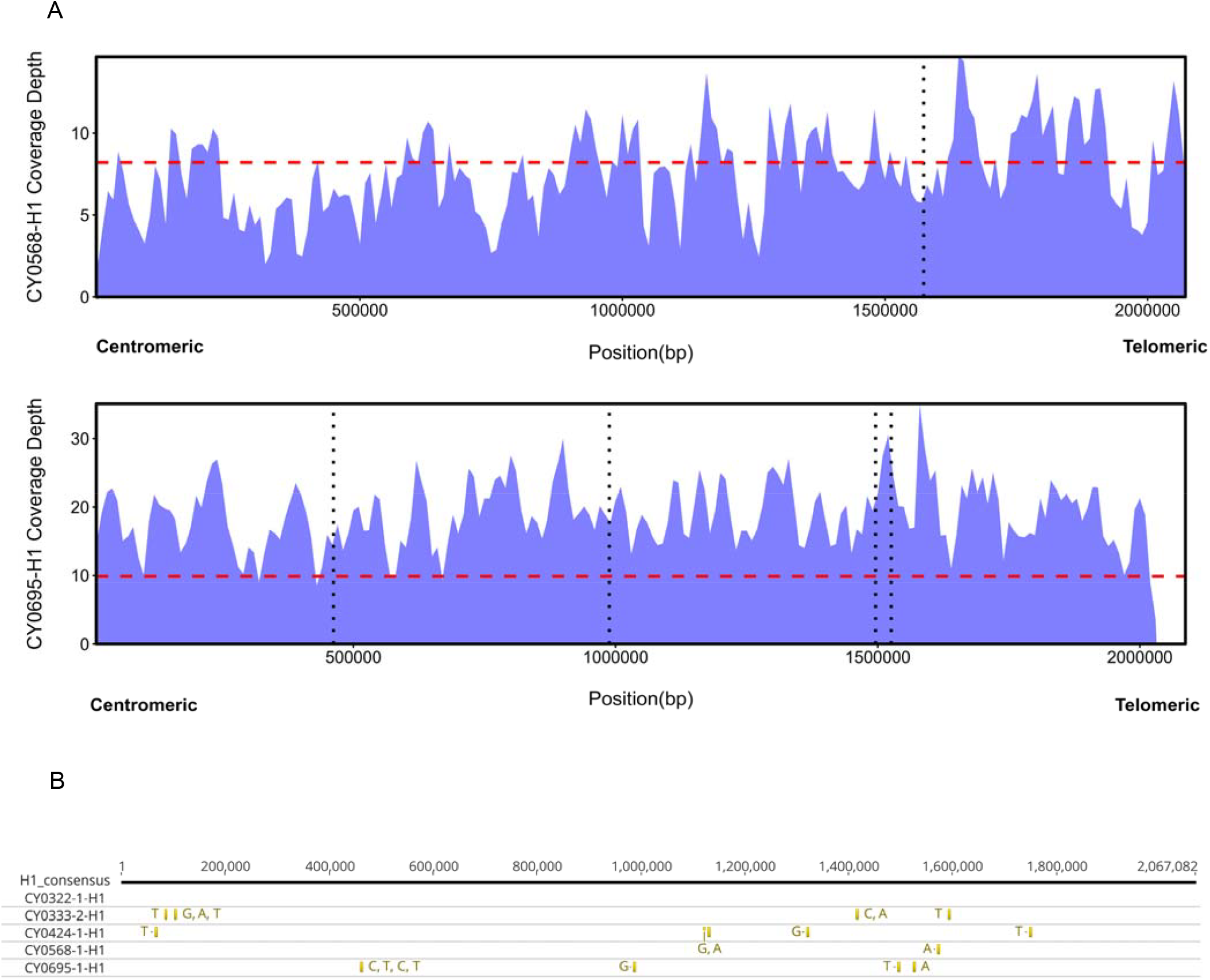
H1 is the predominant IGH haplotype found. (A) Read coverage plots of H1 assemblies from two homozygous animals. Dashed red lines show the average genomic read coverage for each animal. Vertical dotted lines indicate the positions of SNVs shown in panel B. (B) Five full-length, contiguous H1 assemblies differ at a total of just 20 sites across 2 megabases of sequence.

The most common haplotype (“H1”) was present in two of the homozygotes (CY0568 and CY0695) with relatively uniform read coverage and an average depth of 7.4x and 10.3x, respectively, across the IGH locus (Fig 1a). Large spikes in coverage occurred over repetitive elements due to off-target reads mapping to these regions. We identified five additional H1 haplotypes from heterozygous animals based on average sequence identity >99% and no structural variants greater than 2 Kbp. Two of these heterozygous H1 haplotypes in subjects CY330 and CY0558 were fragmented into two and three contigs, respectively (Table 1). We combined the three contiguous H1 haplotypes from heterozygotes with the two homozygous haplotypes to generate an H1 consensus for further analysis (Fig 1b). Remarkably, the five H1 haplotypes were 99.999% identical, differing at only 20 positions across the 2 megabase (Mbp) H1 haplotype. All variable positions fell within intergenic regions (Fig 1, Fig S2). Overall, this suggests that the H1 consensus is representative of the common H1 haplotype and can serve as a reliable genetic reference for analysis of MCM IGH. In addition, haplotypes H2 and H3 were identified from additional homozygous MCM, and haplotypes H4, H5, and H6 were identified from heterozygous MCM (Table 1).

We used ALIGaToR [54] to annotate the IG genes present in H1 (Fig 2a-b). As per the current recommendations of the T-cell Receptor and Immunoglobulin Nomenclature Sub-Committee (TR-IG NSC) of the International Union of Immunological Societies [55], we have assigned temporary labels to these genes using IgLabel [44] (see Methods). Correspondences between temporary labels and the names currently used by KIMDB are available in Table S1. We found 80 functional, 3 ORF, and 40 pseudo IGHV genes; 38 functional and 6 ORF IGHD genes; 5 functional and 2 ORF IGHJ genes; and 8 functional IGHC genes. Of the functional and ORF alleles, 61 IGHV, 29 IGHD, and 7 IGHJ alleles (73%, 66%, and 100%, respectively) were previously annotated in KIMDB [26]. The novel alleles were distributed throughout the locus and across IGHV and IGHD gene families (Fig 2c). Notably, when compared to the IGHV alleles present in the existing IMGT cynomolgus genomic reference, only 69 (83%) of the H1 IGHV alleles had close orthologs (≥95% sequence identity; Fig S3), highlighting the need for updated allele databases fully capture the immunogenetic diversity of this species.

**Figure 2.**
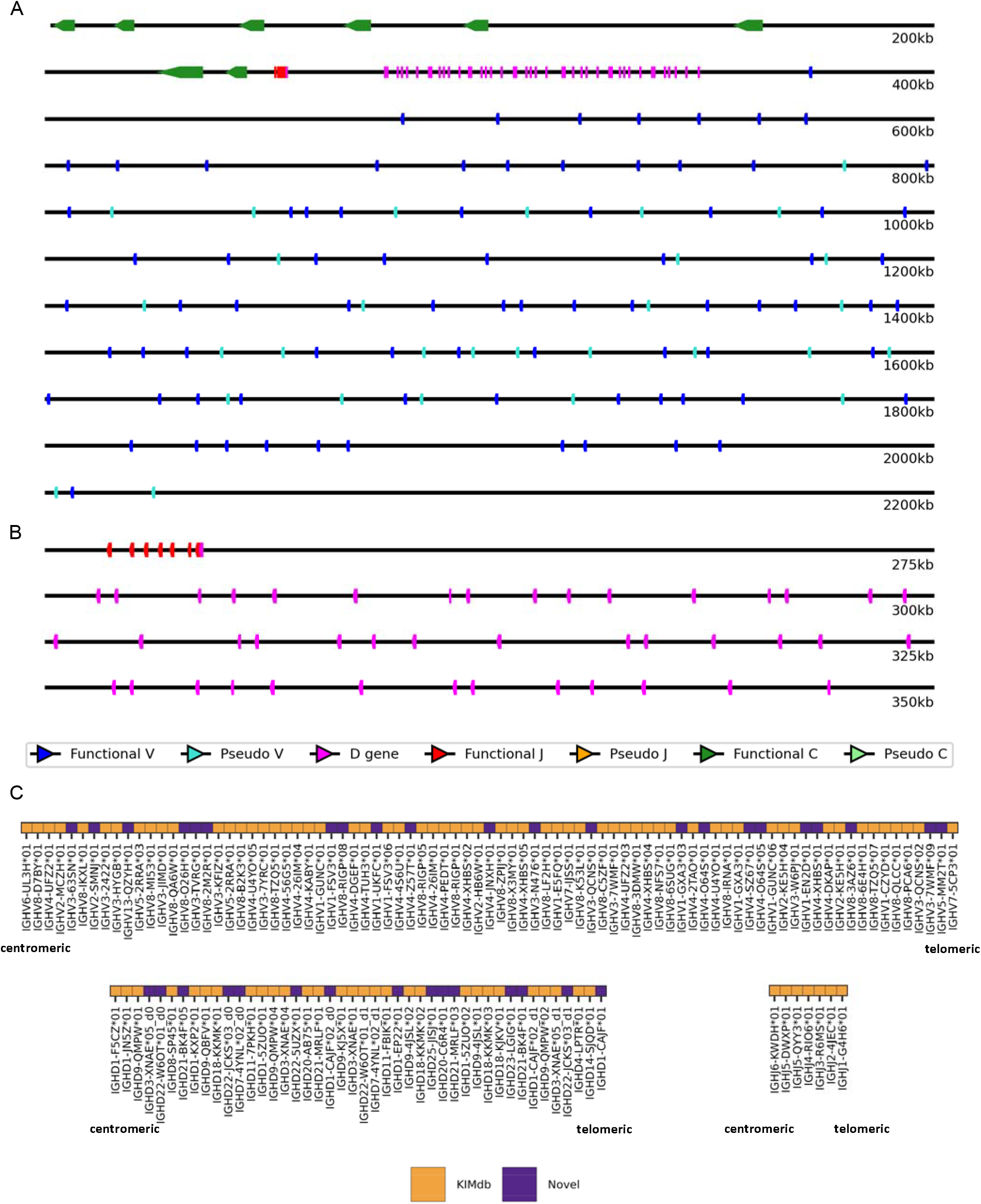
IGH haplotype H1 gene content. (A) Schematic representation of H1 showing the genomic positions of genes annotated by ALIGaToR. (B) Enlarged schematic view of the D-J cluster. (C) H1 V, D, and J genes annotated as functional colored by whether they correspond to alleles represented in the existing KIMdb database. Correspondences between temporary labels and names in KIMdb can be found in Table S1.

### Differences between MCM and the rhesus Mmul_10 genomic reference

We conducted a comparative analysis of the H1 haplotype with the IGH locus of the rhesus macaque reference assembly Mmul_10 [29]. The Mmul_10 IGH locus contains 79 IGHV genes, 44 IGHD, 7 IGHJ, and 8 IGHC genes [56]. The H1 haplotype has one more IGHV genes, six fewer IGHD genes, the same number of IGHC genes and the same number of IGHJ genes though two are designated as ORFs by ALIGaToR (Fig 3). Only 5 IGHV alleles are shared exactly between Mmul_10 and H1, while 7/75 (9%) Mmul_10 IGHV alleles and 11/83 (13%) H1 IGHV alleles share less than 95% sequence identity with any allele in the other haplotype (Fig S4). In addition, there are large structural variations between these haplotypes, consistent with comparisons between other primate species [28]. These structural variations are localized to the IGHV region of the locus and include the insertion or deletion of multiple functional genes, resulting in the H1 haplotype being 103 Kbp larger than the Mmul_10 haplotype. There are also large differences in nucleotide sequence between the H1 and Mmul_10 haplotypes with only 83% identity even in regions that can be aligned between species. Divergence time between rhesus and cynomolgus macaque is estimated at 0.49 to 0.90 million years ago [57]. To compare divergence between rhesus and cynomolgus macaque we aligned whole genome contigs from CY0695 (H1) to Mmul_10 and computed the frequency of single nucleotide variant (SNV) events across all Mmul_10 covered bases. The average frequency of SNV events across the genome is 4.93 bases per 1 Kbp while in IGH only the average frequency is 12.38 bases per 1 Kbp. These results suggest the accelerated divergence of the IGH locus between rhesus macaque and MCM relative to other genomic regions [58] which is consistent with the IGH region in other ape species [28].

**Figure 3.**
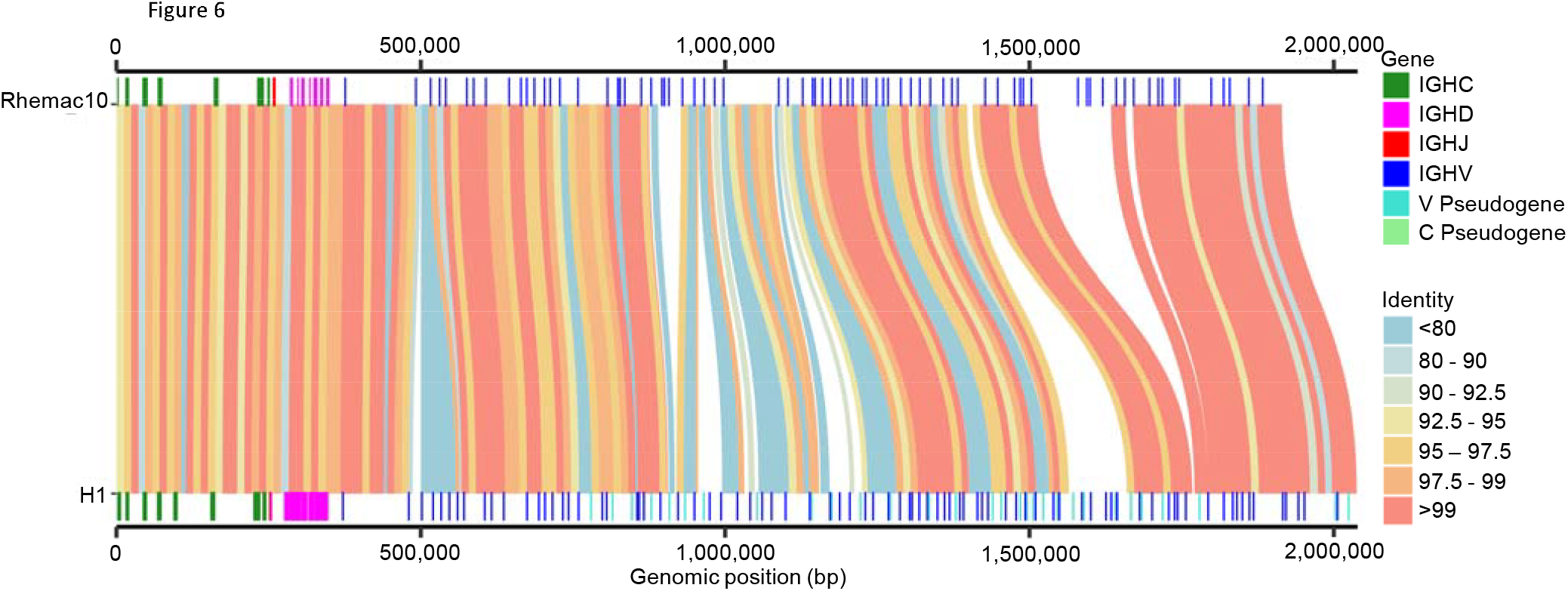
Comparison of IGH from MCM haplotype H1 to M_mul10 reference genome. H1 has approximately 100 kilobases of sequence not found in M_mul10, and overall sequence homology is only 83%.

### Repetitive content within MCM IGH

To evaluate highly repetitive regions across H1 we used the *high multiplicity repeat content* (HMRC) metric to compute the fraction of positions that are covered by repeats of minimum length L at least N times [59]. Positions of repeats were computed by self-alignment of H1 using YASS [60] with a minimum repeat length of 15 Kbp and a minimum coverage of 5. Previous studies have shown that HMRC in the IGH locus can vary significantly across species [59] [28,61]. In H1 we found five HMRC regions in total covering 367 Kbp (18%) of the locus (Fig 4a). Four of the HMRC regions cover a total of 14 V genes (17%) (Fig 4c-d) and a single HMRC region covers 38 (100%) of D genes (Fig 4b). High repeat density across D genes is present in other ape species and human [28,62]. The four HMRC regions across V genes fell in two larger repetitive clusters (Fig 4c-d). The proximal region (Fig 4c) spans ∼450 Kbp, contains 24 functional IGHV alleles and two HMRC regions. The two HMRC regions span 44 Kbp and 30 Kbp respectively; the first contains one IGHV gene and one IGHV pseudogene, while the second contains two IGHV genes which differ from each other by 19 bases (93.6%) and 25 bases (95.8%) in the V exon. The distal region (Fig 4d) is ∼400 Kbp and contains 20 functional IGHV genes and two HMRC regions. The first HMRC region is 155 Kbp and contains eight IGHV genes while the second is 69 Kbp and contains three IGHV genes. These patterns of highly homologous sequence clusters are a common feature of the immunoglobulin loci across different species and are frequently hotspots of structural variation within a species [28,35,63,64]. This demonstrates the power and necessity of long-read sequencing approaches like the one used here, as short-read technologies often fail at resolving complex and highly repetitive regions like the IGH locus [30,65].

**Figure 4.**
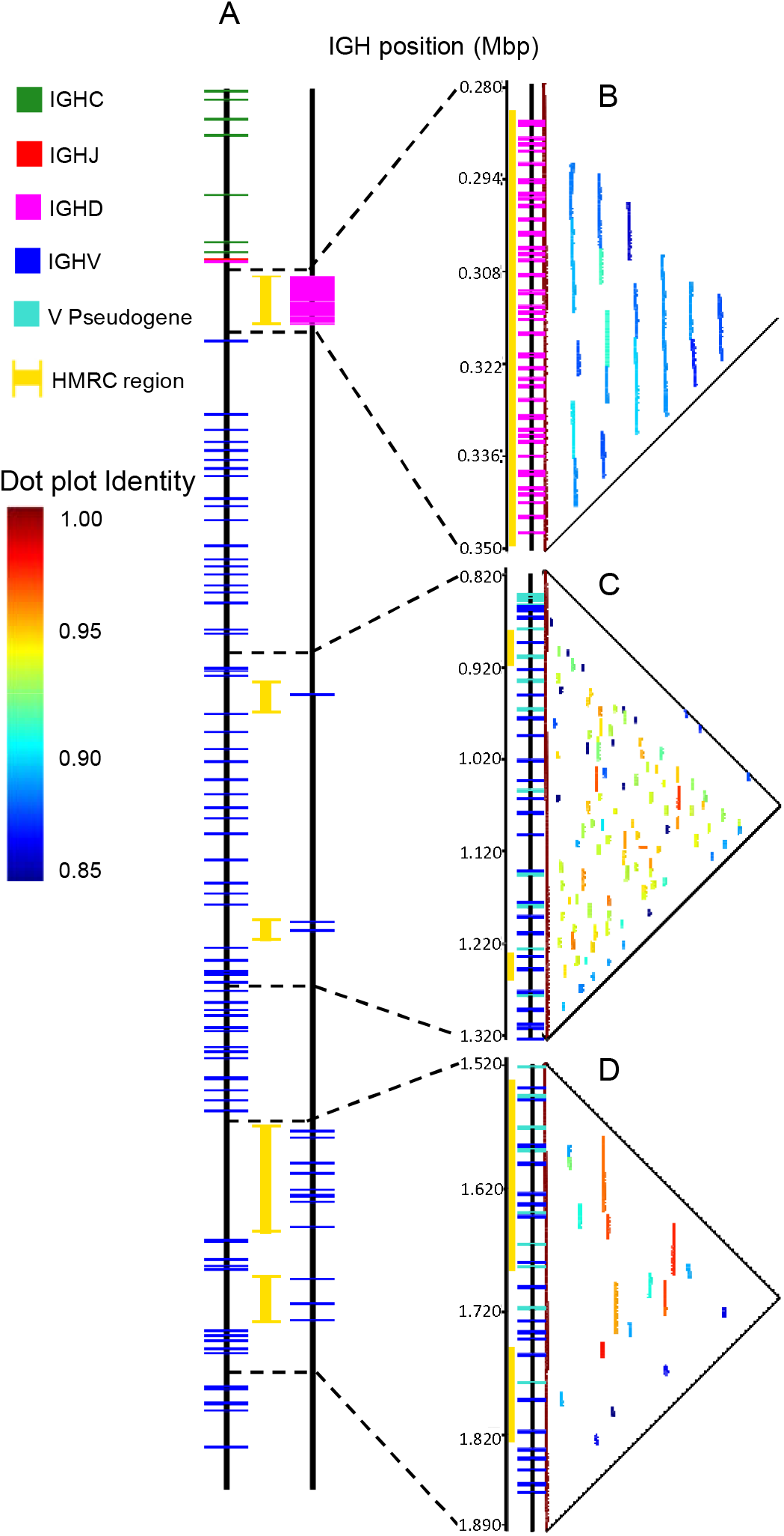
Regions enriched for duplications/repeats within H1. (A) Locus track of the H1 IGH locus. The left-most track are all functional genes outside of HMRC (high multiplicity repeat content) regions (yellow). The right-most track are all genes within HMRC regions. (B) Dot plot of the HMRC region spanning all 38 functional IGHD genes. The dot plot colors correspond to sequence identity. (C) Dot plot of the proximal repetitive region containing two HMRC blocks. The dot plot covers 24 functional IGHV alleles in this region with 3 inside HMRC regions highlighted by a yellow bar. (D) Dot plot of a more distal region with larger homologous blocks and HMRC regions. The dot plot covers 20 functional IGHV alleles in this region with 11 inside HMRC regions highlighted by a yellow bar.

### Structural variation between MCM IGH haplotypes

We then compared the H1 haplotype to two additional haplotypes, H2 and H3, resolved from homozygous MCM. H2 was distributed over 3 scaffolded contigs, but spanned the entire locus, while H3 was comprised of a single full-length contig. Similar to other studies of immunogenetic regions in these animals [16,20], we found a high degree of haplotype diversity (Fig 5). The H1 IGH haplotype is the shortest at 2.07 Mbp, while the H2 and H3 haplotypes are 2.3 Mbp and 2.48 Mbp respectively. The sequence disparity between haplotypes is primarily due to large gene-rich structural variations. The IGHC, IGHJ, and IGHD regions contain little or no structural variation though the sequence identity in these regions varies from ∼92% to >99%. In contrast, the IGHV region contains significant structural variation which primarily occur in or directly adjacent to highly repetitive gene-rich regions. Haplotypes H1 and H2 primarily differ at the proximal end of the IGHV region with interspersed regions of 95-97.5% sequence identity and two large structural variations approximately 600 Kbp and 1,400 Kbp into the locus (Fig 5). The distal end of H1 and H2 have large blocks of highly homologous sequence with >99% identity (Fig 5). Comparatively, the H3 haplotype has a high degree of sequence variation across the entirety of the IGHV region relative to both H1 and H2 (Fig 5). Among regions of structural similarity across the IGH locus, the H1 haplotype has a sequence identity of 99.06% and 98.14% to H2 and H3, respectively.

**Figure 5.**
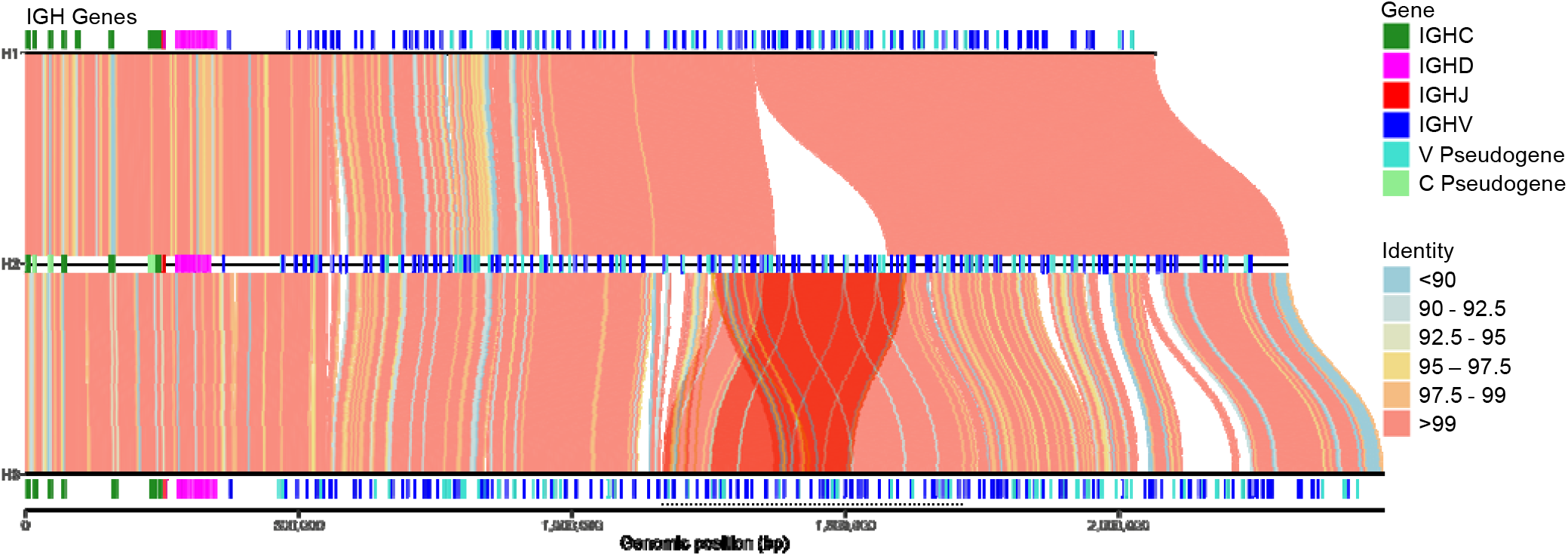
Significant diversity within MCM IGH haplotypes. Alignments of H1, H2, and H3 visualized using SVbyEye. Little structural variation is observed in the IGHC and IGHJ regions (left) with more complex events interspersed with regions of high homology. The underlined region of the H3 haplotype represents a segmental duplication relative to H2.

Relative to H1, the H3 haplotype contains a ∼60 Kbp deletion and a large ∼400 Kbp insertion characterized by multiple segmental duplications in the proximal highly repetitive region (Fig 6a). While the segmental duplications in this region share greater than 99.99% identity, we were able to identify multiple ONT reads that spanned the segmental duplication breakpoints. In this region, H1 contains 26 functional IGHV genes while H3 contains 52. Among these, H1 includes two genes absent in H3 while H3 contains 14 genes not found in H1, as well as a tandem duplication containing 14 genes (28 total). No genes shared between H1 and H3 have identical sequences. Comparing this region across haplotypes H1, H2, H3, and H6 (see below) reveal a potential progression of deletions and duplications that contribute to the divergence between H1 and H3 (Fig 6b). The H2 haplotype retains all 23 genes found in this segment of H1 and includes an additional ∼200 Kbp insertion containing 14 genes that match those in H3. All genes shared between H1 and H2 are sequence identical, with overlapping regions having ≥99.85% sequence identity. Haplotype H6, like H2, contains the ∼200 Kbp insertion absent in H1 but also features a ∼60 Kbp deletion that contains two genes in H1. Of the 38 genes overlapping between H6 and H2, only 10 sequences are identical, and all lie within the ∼200 Kbp insertion region. The H3 haplotype contains an additional ∼200 Kbp duplication that significantly overlaps the insertion found in H2 and H6. Notably, H3 and H6 have 99.88% sequence identity in this region and all 39 distinct genes have identical sequence. In fact, the coding alleles of H3 and H6 are identical, outside of the deletion. This suggests a recent duplication, similar to that observed for the MHC class I *Maga-AG*/*Mafa-G* region in these animals [16]. The H3 and H6 haplotypes are 99.19% identical to H2 and 99.03% identical to H1 with the lowest sequence identity found in the regions immediately flanking the 60 Kbp deletion. Structural variations in the IGH region of humans [35–37] and mice [38,66,67] can significantly impact IGHV gene usage and V(D)J rearrangement. These results suggest that MCMs would be an ideal NHP model to test the impact of structural variation on V(D)J rearrangement and downstream response in therapeutic and vaccine studies, eg by investigating difference in the expressed repertoires between H3- and H6-homozygous animals.

**Figure 6.**
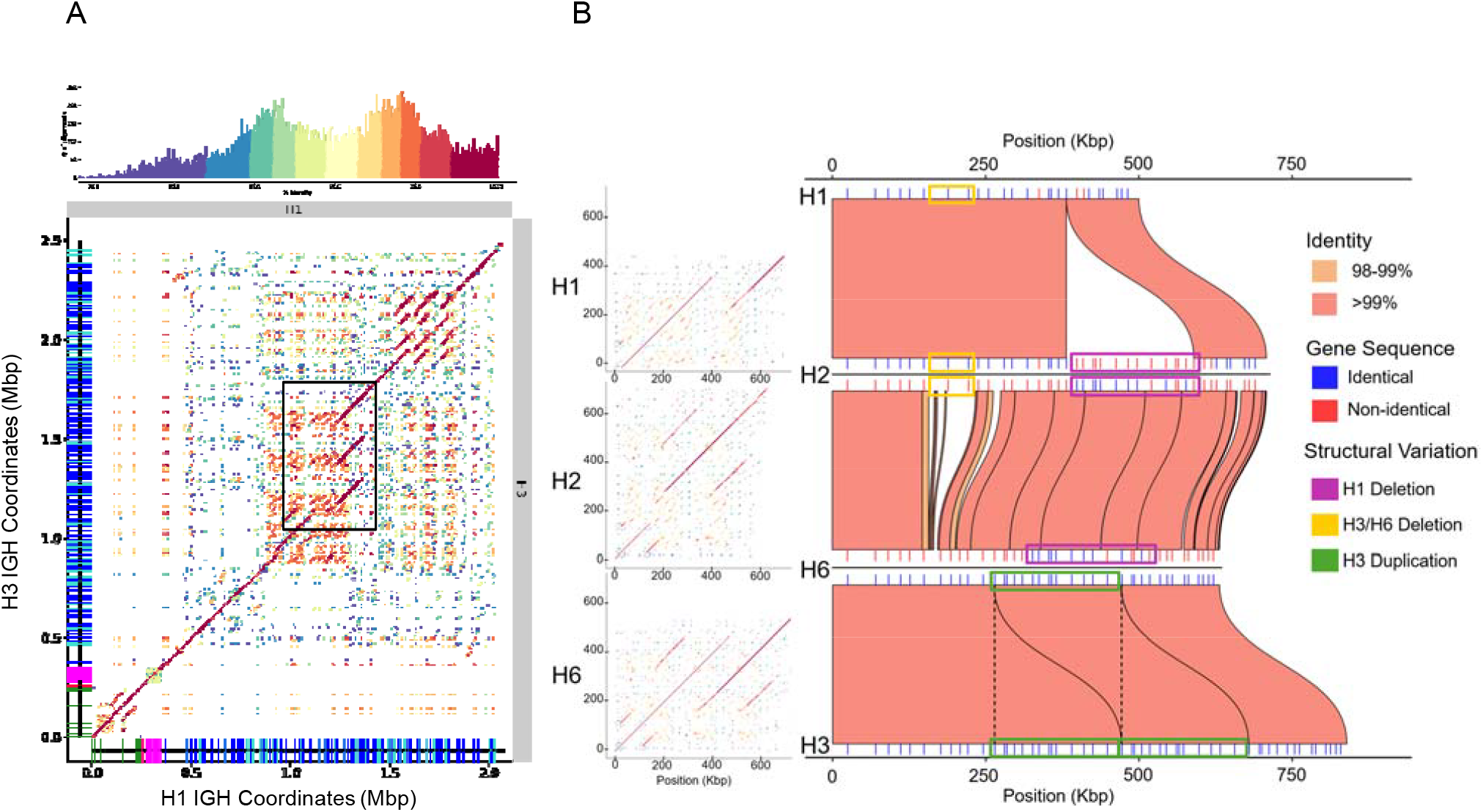
Haplotype H3 contains a large duplications compared to H1. (A) Dot plot comparing H1 (horizontal axis) to H3 (vertical axis). Box (H1:950,000-1,450,000 bp) highlights a region of segmental duplication in H3, just distal of the H1 repeat region show in Fig 3a. The histogram shows the frequency of bins at each identity level, with colors corresponding to colors in the dot plot. (B) Dot plots (left) and Miro plot (right) comparisons of highlighted segmental duplication in haplotypes H1, H2, H3, H6. From top to bottom; H1 is compared to H2, H2 is compared to H6, and H6 is compared to H3. For Dot plots, the top comparison is on the Y-axis and the bottom comparison is on the X-axis. For Miro plots, genes for each haplotype are marked by lines and colored based on sequence identity to comparison haplotype. Blocks of structural variation are highlighted with dotted lines demarcating region of duplication in H3.

### Mapping the range of IGH genetic diversity within MCM

In addition to the complete H1 and H3 haplotypes recovered from homozygous animals, we also recovered a full-length H2 assembly and three additional fully contiguous haplotypes (H4, H5, and H6) from four heterozygous animals, each of which representing a unique haplotype (Table 1, Fig S5). We found that plurality of IGHV alleles (123/288, 43%) are uniquely found in a single haplotype (Fig 7a). An additional 113 alleles (39%) were found in exactly two haplotypes, due to the large blocks of high sequence identity in the IGHV region shared between H1/H2 and H3/H6 (Fig 5, Fig 6b, Fig S6). By contrast, IGHD alleles were predominately seen across all six haplotypes (18/68, 26%) or unique to a haplotype (20/68, 29%). (Fig 7b). No clear pattern was observed for IGHJ alleles (Fig 7c).

**Figure 7.**
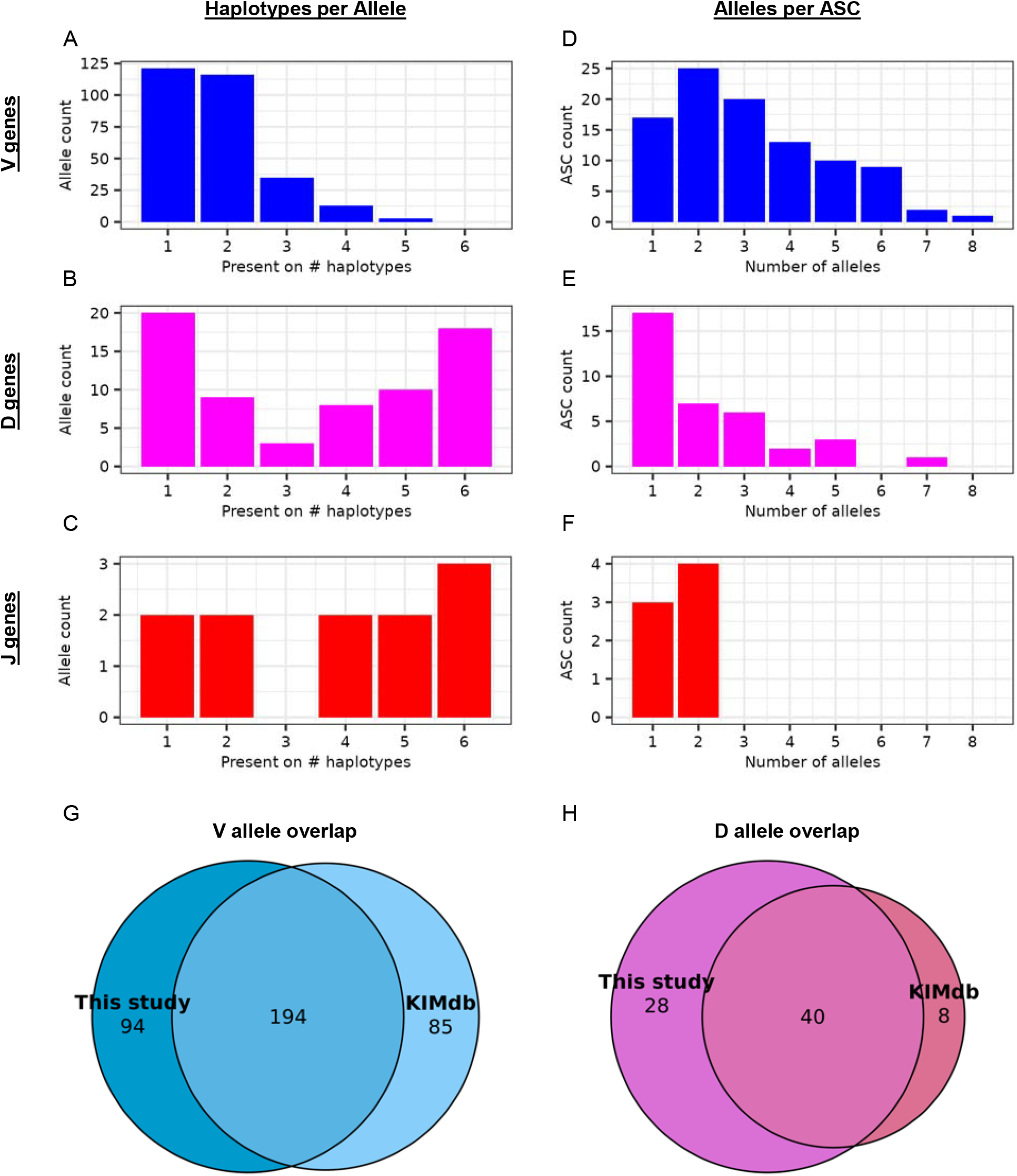
Genetic diversity among MCM haplotypes. (A) The plurality of IGHV alleles appear uniquely in a single haplotype, while (B) IGHD and especially (C) IGHJ alleles are more likely to be shared across most or all haplotypes. The large number of IGHV alleles shared between twohaplotypes is due to extended blocks of shared sequence between H1 and H2, as well as between H3 and H6 (see Fig 6b). (D) IGHV genes are the most diverse, with up to eight alleles per ASC observed in this cohort. (E) IGHD genes and (F) IGHJ genes are less allelically diverse. (G) Overlap between the IGHV alleles observed in this cohort and those from a previous cohort [26]. See also Fig S6. (H) A greater proportion of IGHD alleles are unique to this study due to the difficulty of recovering germline IGHD gene sequences from rearranged IGs.

The structural variation among macaques in immune loci like IGH makes it difficult to align genomes with sufficient resolution to identify sets of alleles that originate from the same gene [30,62,68]. Instead, we used PIgLET [31] to group coding sequences into Allele Similarity Clusters (ASCs), using an identity cutoff of 95% (90% for IGHD alleles) to approximate the similarity among alleles of a single gene. We found 97 IGHV, 36 IGHD, and 7 IGHJ ACSs across all six haplotypes. In general, IGHV ASCs contain multiple alleles per ASC, with 35 IGHV ASCs (36%) having four or more alleles (Fig 7d). In contrast, IGHD ASCs predominantly contained one or two alleles, with only six IGHD ASCs (17%) having four or more alleles (Fig 7e). The IGHJ ACSs followed a similar pattern, with three of the seven ASCs having a single allele with the remainder having two alleles per ASC (Fig 7f). While the shorter length of IGHD and IGHJ alleles compared to IGHV alleles would require a smaller number of nucleotide variants to separate sequences into distinct ASC clusters, the conservation of IGHJ allele sequences and the bimodal frequency of unique and common IGHD alleles across all haplotypes suggest multiple factors influencing the diversity of these genes. Importantly, many IGHD alleles are duplicated, such that there are 68 unique sequences but 81 total alleles. Moreover, some of these duplications are conserved, with all 6 haplotypes carrying at least two identical copies of IGHD22-W6OT*01 and IGHD3-XNAE*05. Notably, IGHD coding alleles appear to be strongly conserved across macaques, with a previous study having found 29/54 (54%) total observed IGHD alleles to be expressed by all animals in all subsets of both Rhesus and Cynomolgus species [26].

To assess the extent of MCM immunogenetic diversity, we compared the alleles recovered from the 13 animals in this study to those inferred from repertoire sequencing of an independent group of 12 MCM [26]. Of 373 total IGHV alleles across both studies, only 194 (52%) were present in both groups of animals (Fig 7g). The 94 alleles that were found exclusively in the current cohort were uniformly distributed across all 6 haplotypes (Fig S6a) and across the locus within each haplotype (Fig 2c). These likely have low usage in the expressed repertoires, which would have prevented their recovery by [26]. To determine if the 85 IGHV alleles present only in KIMDB might represent unmapped haplotypes, we annotated 5 unscaffolded partial IGH contigs (Table 1). This suggested the existence of two additional haplotypes (H7 and H8) and yielded 72 further IGHV alleles, including 41 KIMDB alleles that had not been captured on H1-H6 (Fig S6b). Furthermore, 40/44 remaining KIMDB IGHV alleles not present in the genomic data can be accounted for by pairs of alleles that differ only at position 320 or beyond, which are known to be difficult to infer from rearranged sequences [69] (Fig S6c). The high concordance between the two cohorts provides further evidence that 12 animals are sufficient to capture most of the restricted genetic diversity of MCM even in IGH.

Low overlap was also observed for IGHD alleles, with only 39/76 (51%) observed in both cohorts (Fig 7h), although with significantly more unique alleles in the current study than in the prior cohort. This imbalance likely reflects the difficulty of reconstructing germline IGHD gene sequences from repertoire sequencing data [41,70] rather than true genetic differences between the cohorts. Notably, the addition of the two partial haplotypes did not lead to any further shared alleles between the two databases, consistent with the differences in IGHD not being due to sampling.

We also considered allelic diversity in the constant chains. Eight IGHC genes were annotated in the expected order on all 10 fully spanning assemblies (Table 2). However, IgG4 was not automatically annotated by ALIGaToR in haplotypes H3 and H6 due to loss of the M2 exon. The observation that the H3 and H6 haplotypes share identical allelic variants across the IGHC region suggest that this is not a sequencing or assembly artifact. Further study will be required however, to determine if this represents a functional allele with a truncated intracellular domain or the pseudogenization of IgG4 in these haplotypes.

**Table 2.** IG constant region alleles identified on 10 fully spanning contigs. All eight expected constant alleles were detected in all fully spanning assemblies. Text color indicates the allele number of each gene. For haplotypes H3 and H6, IgG4 was manually annotated (green shading) after failing to be detected by ALIGaToR due to missing M2 exons. Bold font indicates homozygous animals. See also Table 1.

| <u>Animal</u> | <u>Hap</u> | <u>IgM</u> | <u>IgD</u> | <u>IgG1</u> | <u>IgG2</u> | <u>IgG3</u> | <u>IgG4</u> | <u>IgE</u> | <u>IgA</u> |
| --- | --- | --- | --- | --- | --- | --- | --- | --- | --- |
| CY0568 | H1 | IGHM*01 | IGHD*01 | IGHG1*01 | IGHG2*01 | IGHG3*01 | IGHG4*01 | IGHE*01 | IGHA*01 |
| CY0695 | H1 | IGHM*01 | IGHD*01 | IGHG1*01 | IGHG2*01 | IGHG3*01 | IGHG4*01 | IGHE*01 | IGHA*01 |
| CY0322 | H1 | IGHM*01 | IGHD*01 | IGHG1*01 | IGHG2*01 | IGHG3*01 | IGHG4*01 | IGHE*01 | IGHA*01 |
| CY0333 | H1 | IGHM*01 | IGHD*01 | IGHG1*01 | IGHG2*01 | IGHG3*01 | IGHG4*01 | IGHE*01 | IGHA*01 |
| CY0424 | H1 | IGHM*01 | IGHD*01 | IGHG1*01 | IGHG2*01 | IGHG3*01 | IGHG4*01 | IGHE*01 | IGHA*01 |
| CY0161 | H2 | IGHM*01 | IGHD*01 | IGHG1*02 | IGHG2*02 | IGHG3*02 | IGHG4*02 | IGHE*01 | IGHA*01 |
| CY0973 | H3 | IGHM*02 | IGHD*02 | IGHG1*03 | IGHG2*03 | IGHG3*02 | IGH4*05p | IGHE*02 | IGHA*02 |
| CY0161 | H4 | IGHM*03 | IGHD*03 | IGHG1*04 | IGHG2*05 | IGHG3*02 | IGHG4*03 | IGHE*03 | IGHA*03 |
| CY0424 | H5 | IGHM*04 | IGHD*04 | IGHG1*05 | IGHG2*04 | IGHG3*01 | IGHG4*04 | IGHE*04 | IGHA*04 |
| CY0333 | H6 | IGHM*02 | IGHD*02 | IGHG1*03 | IGHG2*03 | IGHG3*02 | IGH4*05p | IGHE*02 | IGHA*02 |

Within IgG, subtypes were labeled by genomic position, rather than sequence homology to other species or inferred function [71]. Similar to previous reports [71] human and macaque IgG clustered separately, with all macaque subtypes being more similar to each other than to any human subtype (Fig 8). Within the macaques, cynomolgus IgG1, IgG2, and IgG4 each clustered together with their rhesus ortholog. Notably, however, IgG3 did not form a distinct cluster, with the MCM alleles instead falling between rhesus IgG3 and cynomolgus IgG4. Additionally, only IGHG3*01, on the H1 and H5 haplotypes, is similar to cynomolgus IgG3 alleles previous documented by IMGT. The other haplotypes carry IGHG3*02, which is in the same relative position as IGHG3*01 but has a shorter hinge region and is closer in sequence to the IGHG4 alleles. Indeed, the coding sequence of IGHG3*02 here is identical to that of cynomolgus IGHG4*02 in the in current IMGT database. It is unclear if these IgGs function more similarly to the cynomolgus IgG4 alleles or the IGHG3*01 allele. Moreover, the diversity in this part of the genome is limited to only 2 total alleles, compared to 4-5 alleles present at each of the other IgG subtype genes. Together, these data suggest a complex evolutionary history in this region.

**Figure 8.**
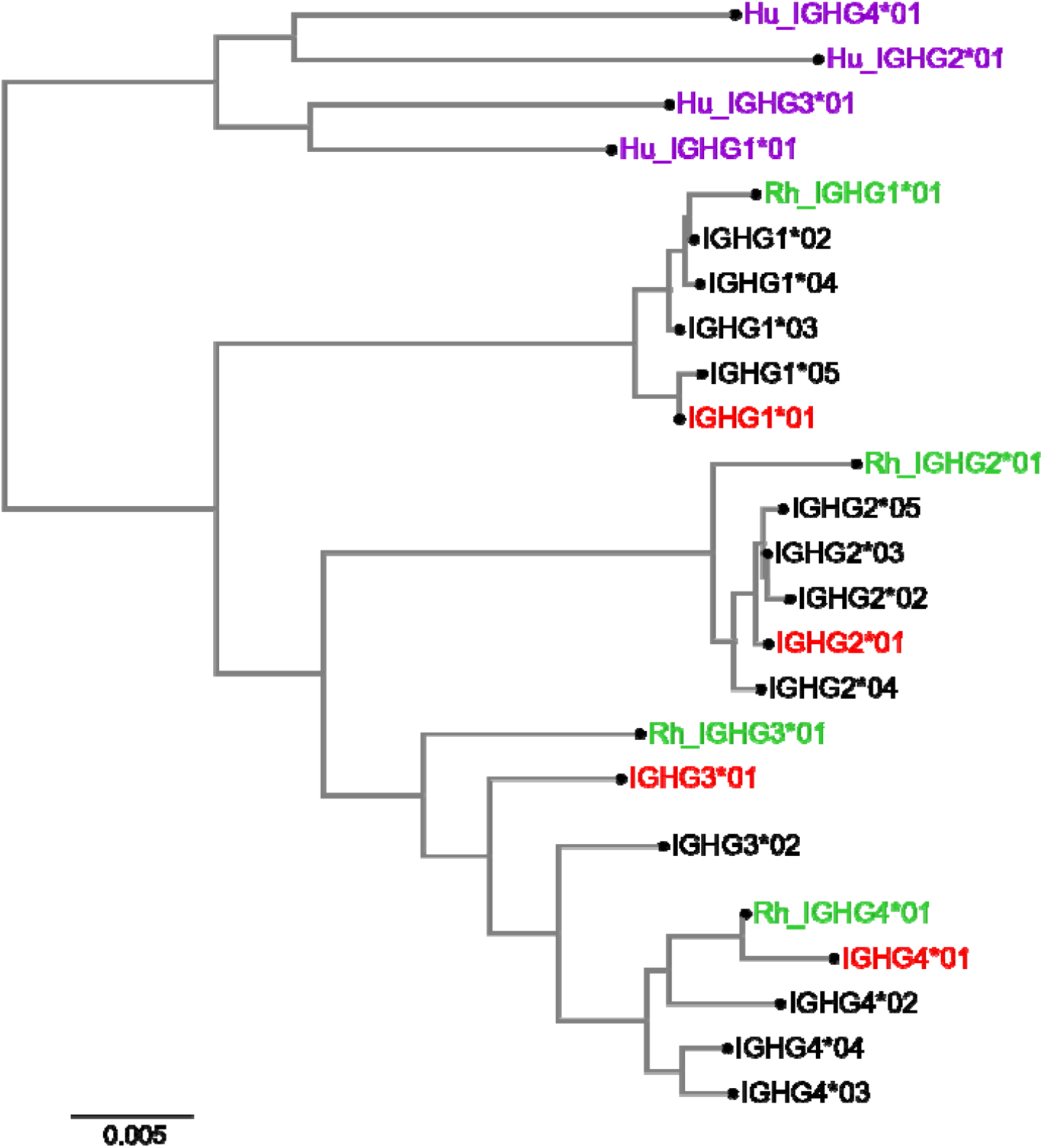
Diversity of MCM IgG constant region subtypes compared to rhesus macaque and humans. Neighbor-joining tree for full-length alleles (CH1 through CHS) of all IgG subtypes found in this study, together with representative alleles from human (purple) and rhesus (green) IgG subtypes. MCM alleles present in H1 are highlighted in red. All other MCM alleles are shown in black.

## Discussion

Overall, we have provided the first full-length, curated genomic reference sequences for IGH in cynomolgus macaques. In addition, by analyzing sequences from 13 animals, we were able to not only characterize inter-species variation between cynomolgus and rhesus macaques, but also assess the immunogenetic diversity within the MCM population, as well. Together with other, immune receptor genes such as MHC, KIR, NKGC, and FCGR, IGs play an outsized role in the course of infectious disease and the response to vaccination [72–76]. Despite this, controlling for genetic variation in these highly polymorphic loci is extremely challenging in outbred species like primates. Crucially, we found that a single dominant haplotype was present in half of the animals studied, highlighting the potential of MCMs for disentangling and assessing the role of genetic factors, due to the restricted genetic diversity in this population.

We have previously used long-read technology to resolve complete haplotypes of the genetic loci for MHC, KIR, natural killer group 2 (NKG2) receptors, and Fcγ receptors in MCM [16,20,76]. Here, we assembled and characterized 6 full-length haplotypes of MCM IGH, including three from homozygous animals. The predominant haplotype, H1, accounted for half of the full-length IGH assemblies recovered. These results are consistent with our previous analyses of these animals, which found that 3 MCM in this study are homozygous for a common NKG2 haplotype (N1) with a frequency of 40% [20]. As these are the first full-length curated assemblies of the IGH region in cynomolgus macaques, this represents a significant advance in the genomic resources available for this species. Moreover, knowledge of the specific IgG alleles carried by MCM will provide a fuller understanding of their interactions and binding affinity to the Fcγ receptors present in this subspecies [76].

Collectively, the haplotypes reported here identify a high total degree of sequence diversity moderated by the relative abundance of a common haplotype. Regions of high sequence homology between haplotypes can nonetheless contain structural variations (Fig 5), with both small (Fig 4) and large (Fig 6) segmental duplications in gene-rich regions. Haplotype diversity is even more extreme when compared to rhesus macaque (Fig 3), further signifying the importance of accurate genomic references to account for genetic variation. Structural variation in the IGH, IGK, and IGL loci of humans has been correlated with differential gene usage [35–37]. While SNV and structural variation in the IGH locus can significantly alter gene usage, the mechanisms and impact of genetic variation on the expressed repertoire is not well understood. The diverse haplotypes characterized from this genetically restricted population represent a powerful model to further explore the associations between germline and phenotype. For example, a vaccine study involving two separate groups of genetically restricted MCMs can elucidate the impact of genetic features on vaccine response.

Our study is limited by the small number of animals sequenced and the limited sequencing depth achieved for some of the animals, especially for the ONT reads necessary to span large SVs and scaffold assemblies of the entire IGH locus. Thus, we fully assembled 13 loci, which, accounting for the 4 homozygous animals, cover 17/26 (65%) copies of the IGH locus. Analysis of partial assemblies and comparisons to an existing database indicated the presence of at least two additional haplotypes beyond the six analyzed here. In addition, 6/10 (60%) of the animals with at least one complete assembly are heterozygous, raising the possibility of assembly errors confounding our analyses [77]. Finally, we currently lack sufficient data to fully trace the history of complex structural variations across haplotypes and confidently link alleles from orthologous genes. Instead, we have relied on ASCs, which collapse evolutionarily recent duplications that still maintain high sequence homology.

In addition to structural variation across haplotypes, more data are needed to distinguish between the likely ancestral IGH haplotypes of the MCM founder population and more recent recombinants. Previous work has identified 7, 8, 8 and 7 MCM haplotypes for MHC, KIR, FCGR and NKG2, respectively [76,78–81]. Each of these loci is on a different chromosome and therefore expected to segregate independently. Similarly, we currently describe 6 IGH haplotypes, with evidence for two additional haplotypes present in these animals (Table 1) and other MCM (Fig 7g, Fig S6b). Moreover, we found large blocks of high sequence homology between H1 and H2 (Fig 5, Fig 6b), as well as between H3 and H6 (Fig 6b). The restricted genetic diversity of this subspecies thus provides a tractable system for the analysis of genetic effects on the expressed repertoire and responses to vaccines and other immunological challenges.

## Conclusions

In total, we annotated over 400 unique alleles across all IGH gene types, 132 (33%) of which are not present in a previous MCM database. Furthermore, 43% of IGHV alleles and 29% of IGHD alleles were unique to a specific haplotype (Fig 7 a-b). The large number of novel alleles and the diversity in gene and allele content between haplotypes demonstrates the necessity of genotyping study animals and continuing to add to databases. Future work will link genomic analysis with AIRR-seq data to better understand the functional effects of genetic diversity[35,45], with the potential to provide insights into the biological mechanisms underpinning different immune responses. Overall, the extensive analysis of the IGH region here has revealed significant biological implications for the usage of MCMs and cynomolgus macaques in biomedical research.

## Methods

### Assembly and mapping

Whole genome sequencing using PacBio and Oxford Nanopore Technology for 13 MCM were retrieved from NCBI (PRJNA854973 [20]) and assembled *de novo* with Hifiasm [52,53] using the command:

~~~
**hifiasm -o {output.asm} -t16 --ul {Nanopore.fq.gz} {PacBio.fq.gz}**
~~~

Contigs either partially or fully spanning the IGH locus were identified using BLAST+ with the Mmul_10 IGH locus as the query. Contigs spanning the centromeric edge of the IGH locus were trimmed using an anchor sequence ∼2 Kbp centromeric of the IgA gene. To assess assembly quality over the IGH locus HiFi and Nanopore reads were mapped back to the assembly contigs and IGH locus contigs were visually inspected [82]:

~~~
**minimap2 -ax map-pb {reference}.fa {reads.fastq} > {output.sam}**
~~~

Coverage was calculated using `samtools coverage` and plotted in R using `ggplot2`.

The H1 consensus was generated in Geneious Prime version 2022.1.1 (Biomatters Ltd.) and minimap2 was used to map each full-length H1 contig against the consensus:

~~~
**minimap2 -ax asm20 {H1-cons.fasta} {input.fasta} > {output.sam}**
~~~

and bcftools [83] was used to call variants:

~~~
**bcftools mpileup -f {H1-cons.fasta} {input.bam}** |
      **bcftools call -mv -o {input.vcf}**
~~~

The SNVs were then visualized in Geneious.

### Gene Annotation

ALIGaToR [54] was used to annotate the IGH assemblies starting from the IMGT [84] annotations of the Mmul_10 reference genome. Recombination signal sequences (RSS) were predicted based on RSS Information Content [85] using Dnagrep [86] in ALIGaToR. *M. fascicularis* IGHV, IGHD, and IGHJ coding sequences were downloaded from KIMDB [26] and filtered to only those identified as present in at least one MCM. These were combined with coding sequences of *M. fasciularis* IGHC from IMGT and used to identify shared alleles. The final ALIGaToR command was:

~~~
**aligator annotate contig.fasta contig.RSS12.bed contig.RSS23.bed \**
      **IGH IMGT_reference.fasta IMGT_reference.bed --outgff contig.gff \**
      **--outfasta functional_genes.fa \**
      **--alleledb KIMdb_IMGT_reference_sequences.fa \**
      **--psfasta pseudo_genes.fa**
~~~

Piglet [31] was then use to separately cluster all unique coding alleles of V, D, and J genes across the 27 IGH contigs (Table 1) and the KIMDB MCM subset. V and J genes used ‘`allele_cluster_threshold=95`’, while D genes were clustered with ‘`allele_cluster_threshold=90`’. For V genes only, ‘`trim_3prime_side=318`’ was used, while D and J had ‘**trim_3prime_side=*NULL***’. For each ASC, IgLabel [44] was used to assign a single temporary label, with allele numbers added manually based on ASC membership. For IGHV and IGHD, family numbers are matched as much as possible to those reported in a recent immunogenetic survey of rhesus macaques [45], though individual ASCs were not screened for sequence homology to that database. For IGHC, allele numbers were assigned manually with *01 corresponding to the H1 haplotype and no screening for homology to alleles in the current IMGT database (of which only 3 were observed in these animals).

The annotations were visualized as a number line schematic using `matplotlib`. Tile plots and bar graphs were visualized using `ggplot2`. Venn diagrams were created using the `eulerr` package in R. The neighbor-joining tree was generated using the `ape` package and visualized using `color.plot.phylo` from the `picante` package.

### Identifying High Multiplicity Repeat Content Regions

To identify regions of HMRC we first self-aligned the H1 haplotype to itself using YASS[60] using default settings and the -d 2 option for tabular blast format output. This table was then filtered to remove alignments less than L=15 Kbp and to remove self-alignments where query and target coordinates are equal. We then calculated the number of unique target alignments over every query region to determine the regions with coverage N=5 or greater and filtered out the rest. Any overlapping regions were then merged. Due to the presence of telomeric sequence at the distal end of the H1 haplotype, 27,752 bp of telomeric sequence was masked from this analysis.

### Structural Variation

Alignments between haplotypes were generated using minimap2 [82]:

~~~
**minimap2 -x asm20 -c -eqx -secondary=no {input_1.fasta} {input_2.fasta} >**
**{output.paf}**
~~~

SVbyEye [87] was used to visualize sequence alignments. A minimum alignment size cutoff of 9000, 13000, and 10000 were used to generate Fig 3, Fig 5, and Fig 6 respectively. To filter out alignments, the `breakPaf` or `breakPafAlignment` commands were used to expand PAF alignments which were then subset to the specified cutoffs. In addition, the SVbyEye commands `addAnnotation` and `plotSelf` were modified to replace default coloring and shape parameters with custom coloring and shape parameters. The locus coordinates used for Fig 6 are H1:950,000-1,450,000, H2:989,287-1,697,206, H6:964,653-1,595,927, and H3:964,529-1,803,724. PatchWorkPlot [88] (https://github.com/yana-safonova/PatchWorkPlot) was used to generate dot plots of self-alignments (Fig 4). The command used was ‘`python PatchWorkPlot.py -i input_config.csv -o patchwork_output --cmap jet -- reverse-cmap false’` with input fastas corresponding to the different haplotypes or regions of focus. StainedGlass [89] was used to generate dot plots of haplotype comparisons (Fig 6). In brief, sequences for comparison were indexed using samtools [83] and used as inputs for the StainedGlass snakemake pipeline using default parameters.

## Declarations

### Ethics approval and consent to participate

Animal selection PBMCs and splenocytes were obtained from 13 MCM housed at the Wisconsin National Primate Research Center during semiannual health checks or at necropsy. The animals were selected for analysis based on previously established MHC and KIR genotypes [78,81,90,91]. Sampling was performed in concordance with protocols approved by the University of Wisconsin-Madison Institutional Animal Care and Use Committee, as well as guidelines contained within the Animal Welfare Act, the Guide for Care and Use of Laboratory Animals, and the Weatherall report [92]

### Consent for publication

Not applicable

### Availability of data and materials

Genomic sequencing for this study was downloaded from NCBI BioProject with accession number PRJNA854973. Assembled IGH contigs and annotations have been deposited in Genbank with accession numbers PX411181-PX411201.

### Authors’ contributions

Conceptualization: RWW, DHO, DCD, CAS

Data curation: SO, WSG, TMP, CAS

Formal Analysis: SO, WSG, CAS

Investigation: TMP, JAK

Supervision: RWW, DHO, DCD, CAS

Visualization: SO, WSG, CAS

Writing – original draft: SO, WSG, CAS

Writing – review & editing: all authors

## Acknowledgements

Not applicable

## Supplemental Information

**Figure S1. Comparison of read coverage for IGH assemblies from homozygous animals**. (A) CY0692 and (B) CY0973 versus (C) both assemblies from heterozygous animal CY0161. The assembly for CY092 is fragmented into 3 contigs. Dashed red lines show average genomic coverage for each animal. and plots, bin size of 10,000bp. Fragmented homozygous haplotype CY0692.

**Figure S2. Read support for each of the 20 H1 SNVs relative to consensus shown in Fig 1b**.

**Figure S3. Current databases do not adequately capture the immunogenetic diversity of cynomolgus macaques**. Heatmap showing nucleotide sequence identity between MCM IGHV alleles observed on haplotype H1 (rows) and known cynomolgus macaque IGHV alleles from IMGT (columns). Although H1 is a commonly observed haplotype in our cohort, only 69/82 (84%) of H1 IGHV alleles have at least 95% sequence identity to at least one allele in the IMGT database.

**Figure S4. Extensive allelic differences between species**. Heatmap showing nucleotide sequence identity between MCM IGHV alleles observed on haplotype H1 (rows) and IGHV alleles observed on the rhesus macaque Mmul_10 reference genome (columns). Only 5 alleles are shared exactly between species and 11/82 (13%) of H1 IGHV alleles have no counterpart in Mmul_10 with at least 95% sequence identity.

**Figure S5. Significant diversity within MCM IGH haplotypes**. Dotplots of H1, H2, H3, H4, H5, and H6 visualized using PatchWorkPlot [88].

**Figure S6. Differences between KIMDB and the current study**. (A) Each haplotype identified in this study includes 21-33 IGHV alleles that were not captured in KIMDB. See also Fig 2c. (B) When alleles from two partial additional haplotypes (Table 1) are included, 41 alleles previously found only in KIMDB are now identified as shared alleles. (C) When differences at or after position 320 are ignored, 40 KIMDB-only alleles collapse onto 38 alleles identified in the current study, including 10 other KIMDB alleles. Together with the two partial haplotypes, this eliminates almost all potential false negatives in the current study. (D) Only 3 additional IGHD alleles are identified in the partial haplotypes, none of which were previously documented in KIMDB.

**Table S1. Sequences of alleles reported with corresponding KIMDB labels**.

