## Supplemental Figures 1-6 for "Multiple full-length homozygous IGH haplotypes from Mauritian cynomolgus macaques"

### Slide 1
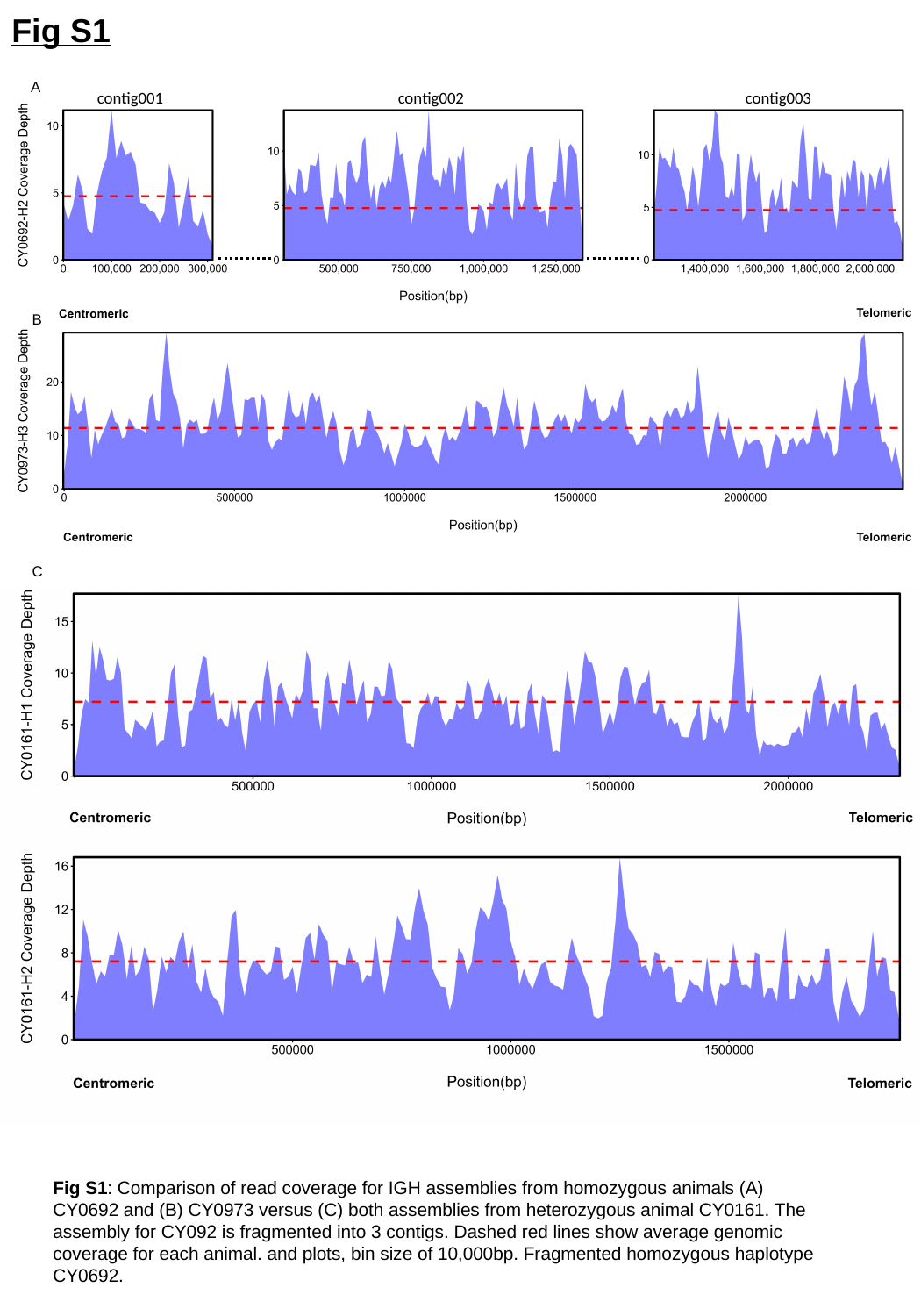

Fig S1
A
contig002
contig003
contig001
B
C
Fig S1: Comparison of read coverage for IGH assemblies from homozygous animals (A) CY0692 and (B) CY0973 versus (C) both assemblies from heterozygous animal CY0161. The assembly for CY092 is fragmented into 3 contigs. Dashed red lines show average genomic coverage for each animal. and plots, bin size of 10,000bp. Fragmented homozygous haplotype CY0692.

### Slide 2
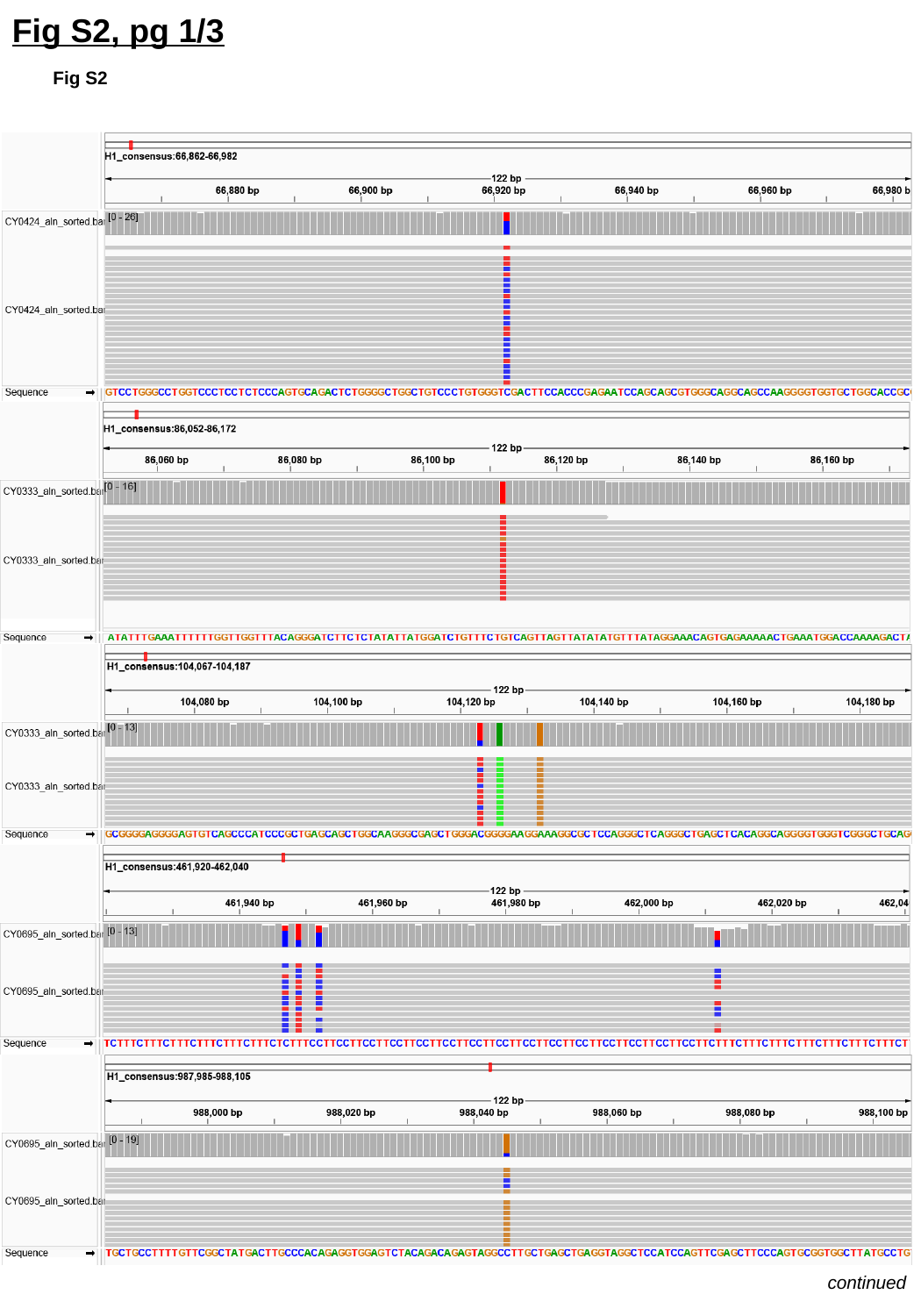

Fig S2, pg 1/3
Fig S2
continued

### Slide 3
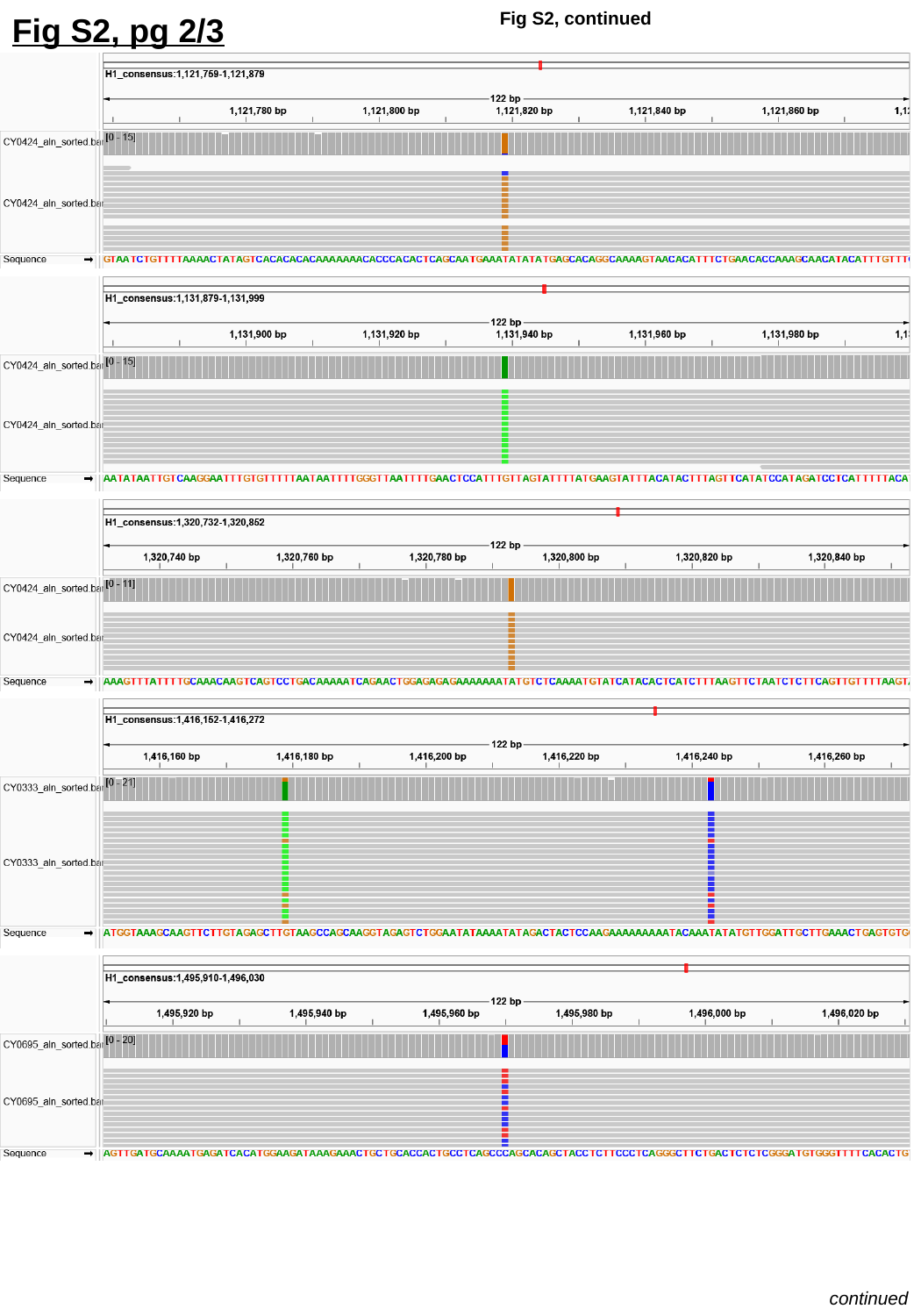

Fig S2, continued
Fig S2, pg 2/3
continued

### Slide 4
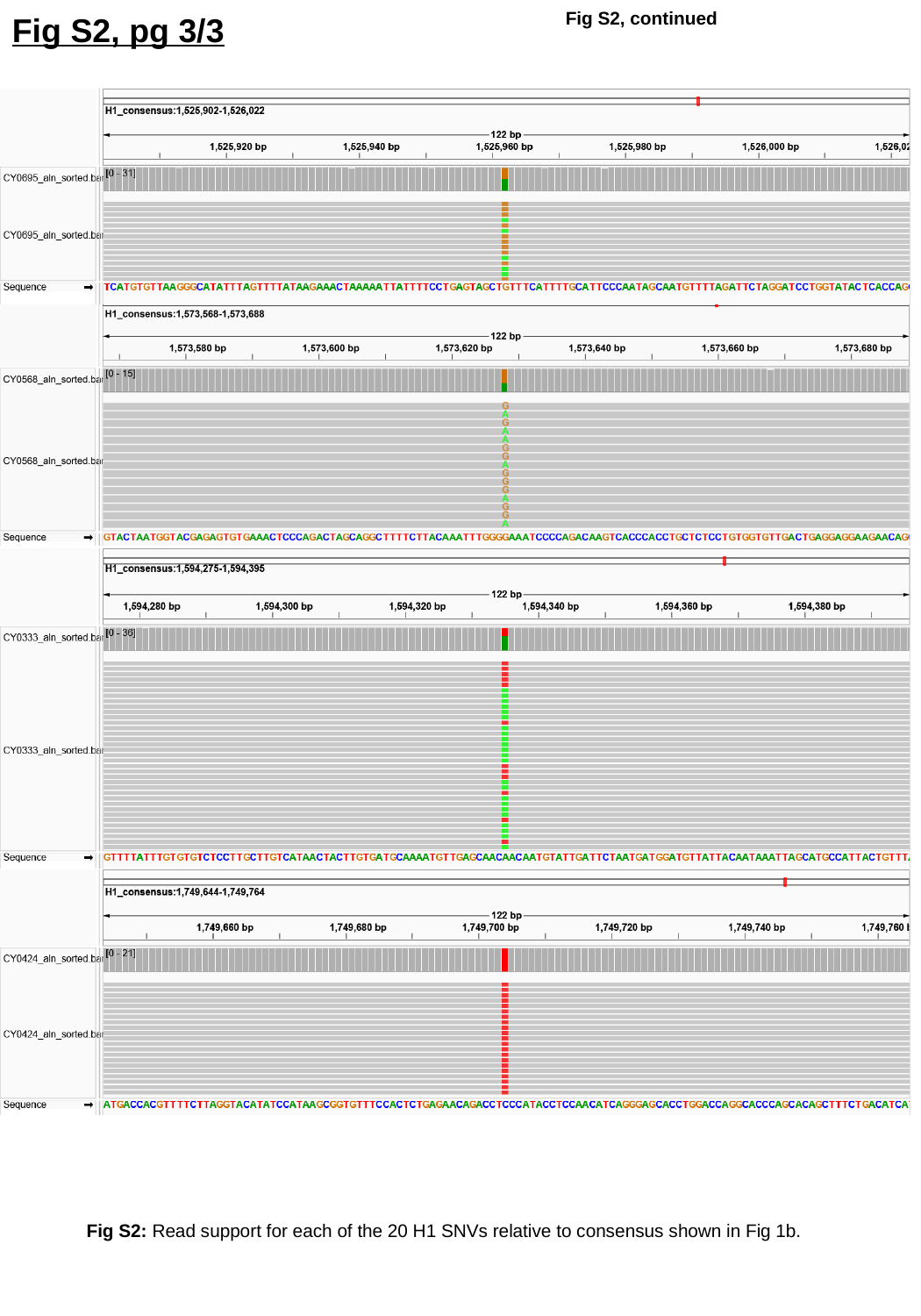

Fig S2, continued
Fig S2, pg 3/3
Fig S2: Read support for each of the 20 H1 SNVs relative to consensus shown in Fig 1b.

### Slide 5
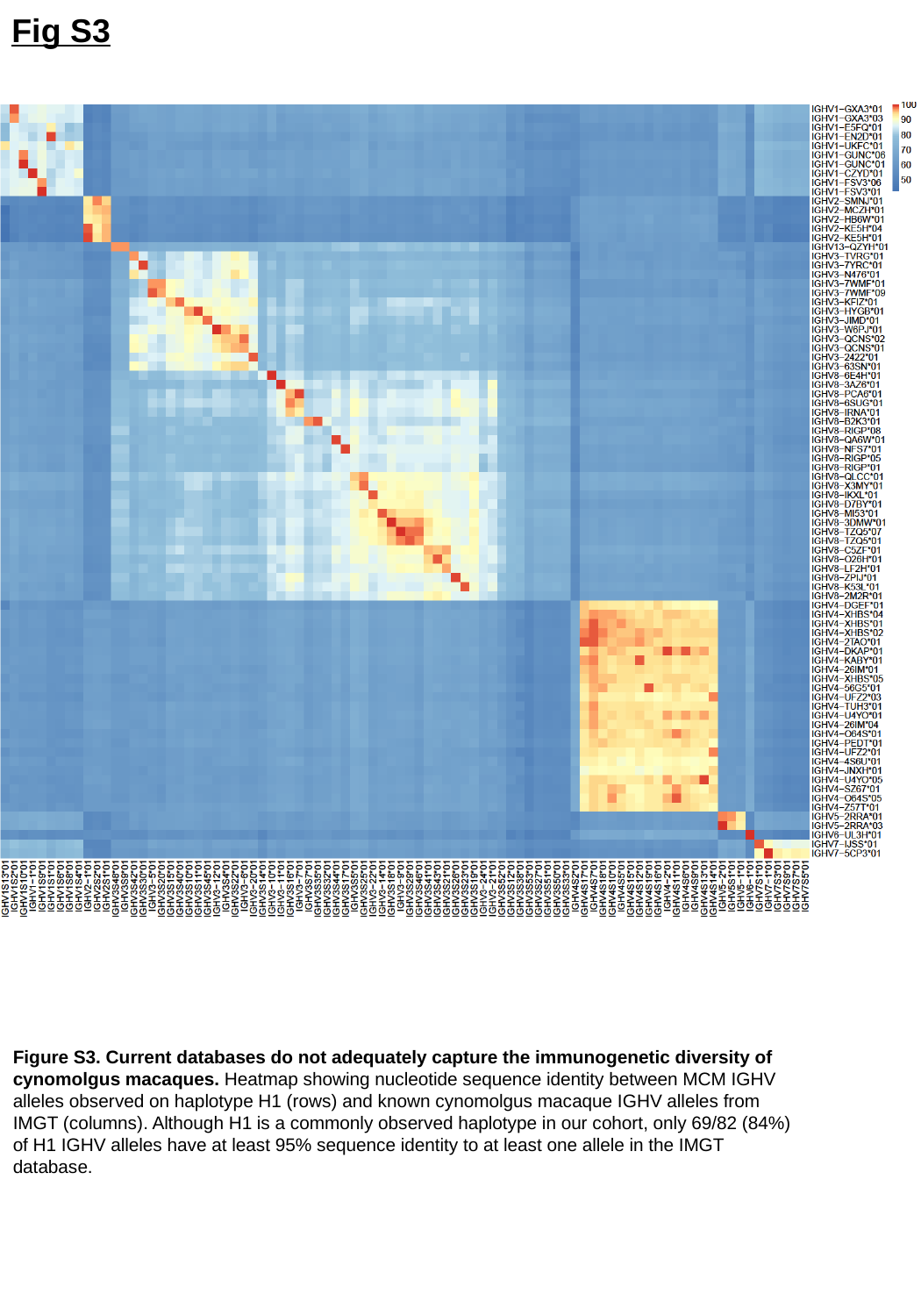

Fig S3
Figure S3. Current databases do not adequately capture the immunogenetic diversity of cynomolgus macaques. Heatmap showing nucleotide sequence identity between MCM IGHV alleles observed on haplotype H1 (rows) and known cynomolgus macaque IGHV alleles from IMGT (columns). Although H1 is a commonly observed haplotype in our cohort, only 69/82 (84%) of H1 IGHV alleles have at least 95% sequence identity to at least one allele in the IMGT database.

### Slide 6
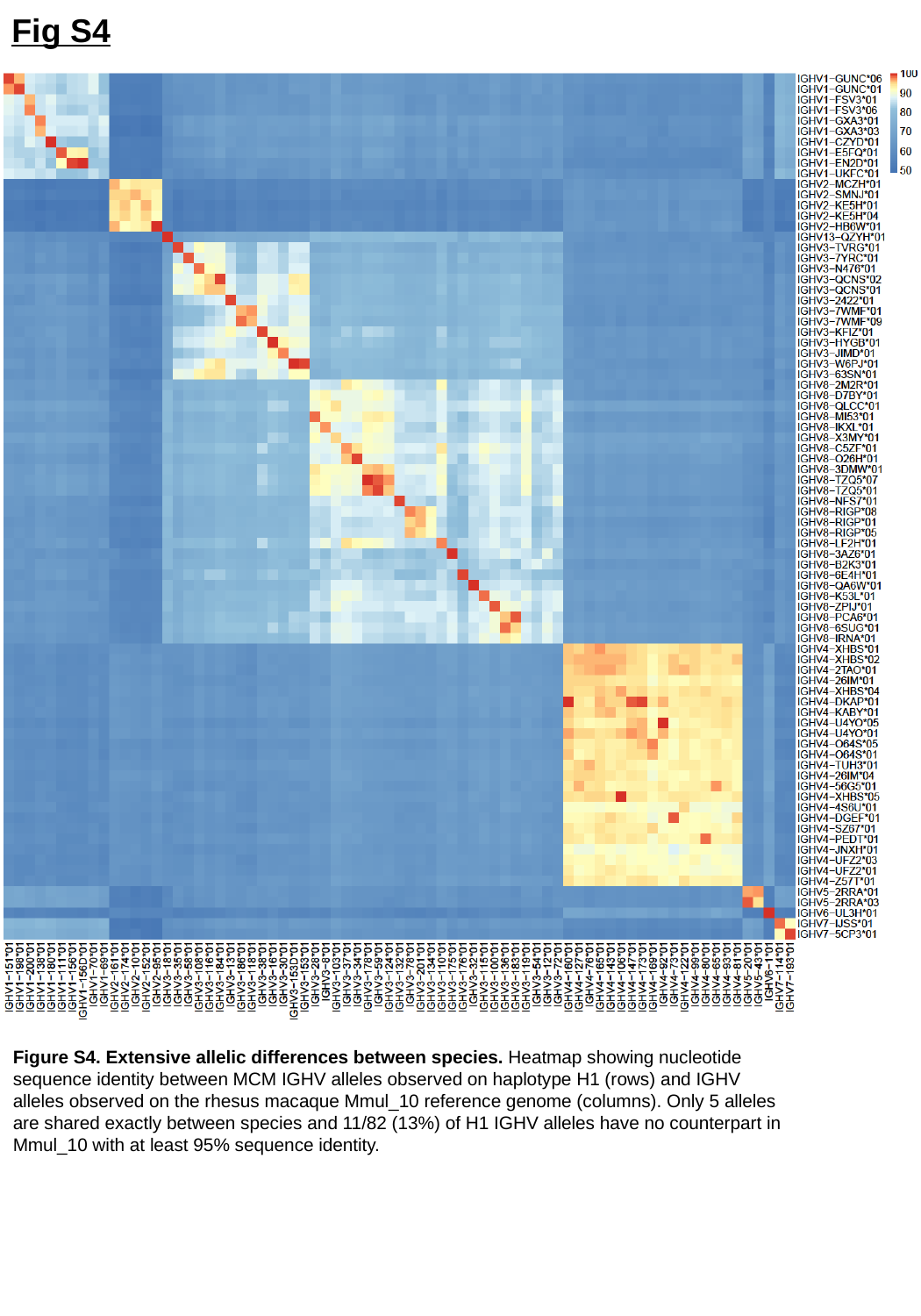

Fig S4
Figure S4. Extensive allelic differences between species. Heatmap showing nucleotide sequence identity between MCM IGHV alleles observed on haplotype H1 (rows) and IGHV alleles observed on the rhesus macaque Mmul_10 reference genome (columns). Only 5 alleles are shared exactly between species and 11/82 (13%) of H1 IGHV alleles have no counterpart in Mmul_10 with at least 95% sequence identity.

### Slide 7
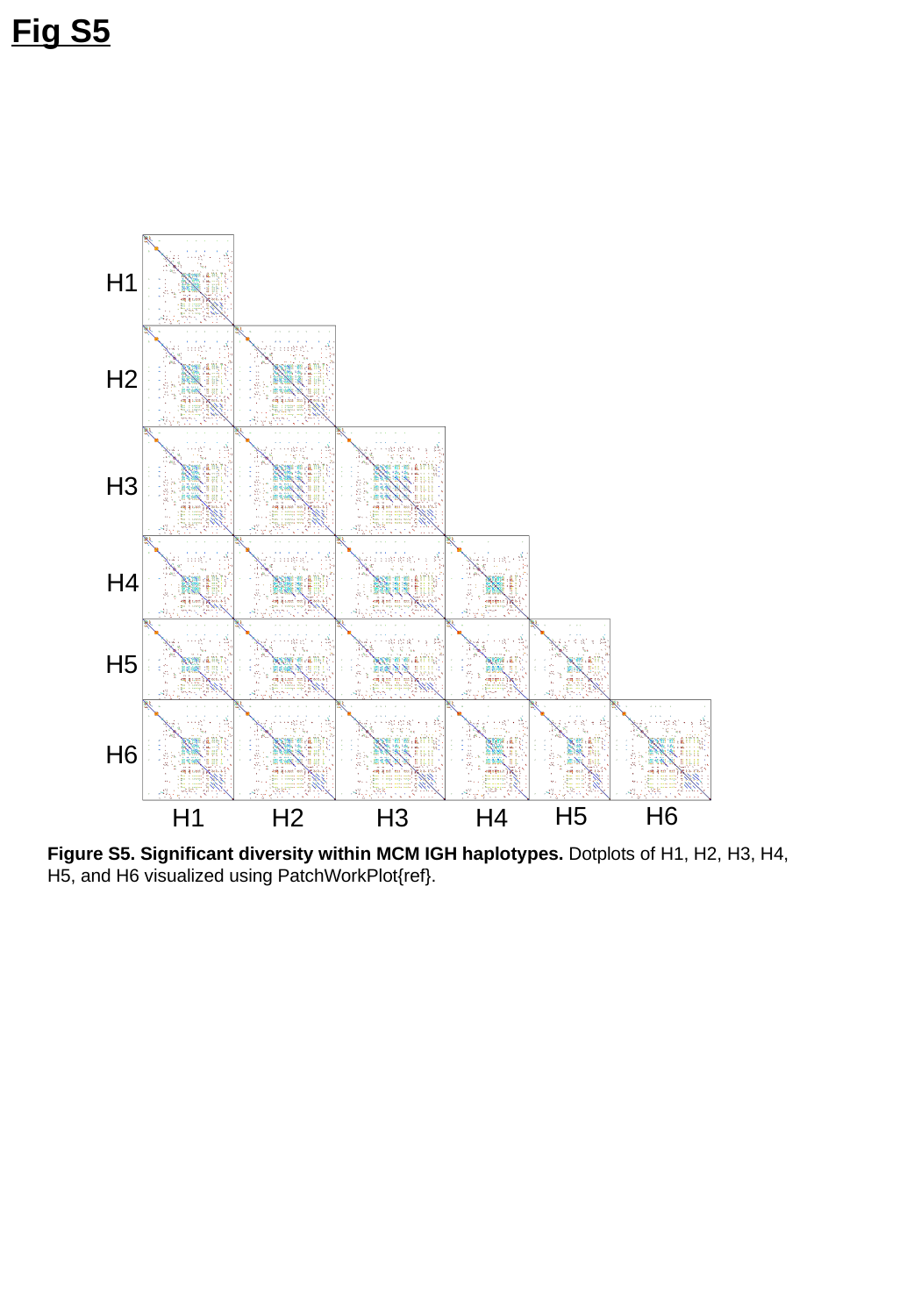

Fig S5
H1
H2
H3
H4
H5
H6
H6
H5
H2
H4
H1
H3
Figure S5. Significant diversity within MCM IGH haplotypes. Dotplots of H1, H2, H3, H4, H5, and H6 visualized using PatchWorkPlot{ref}.

### Slide 8
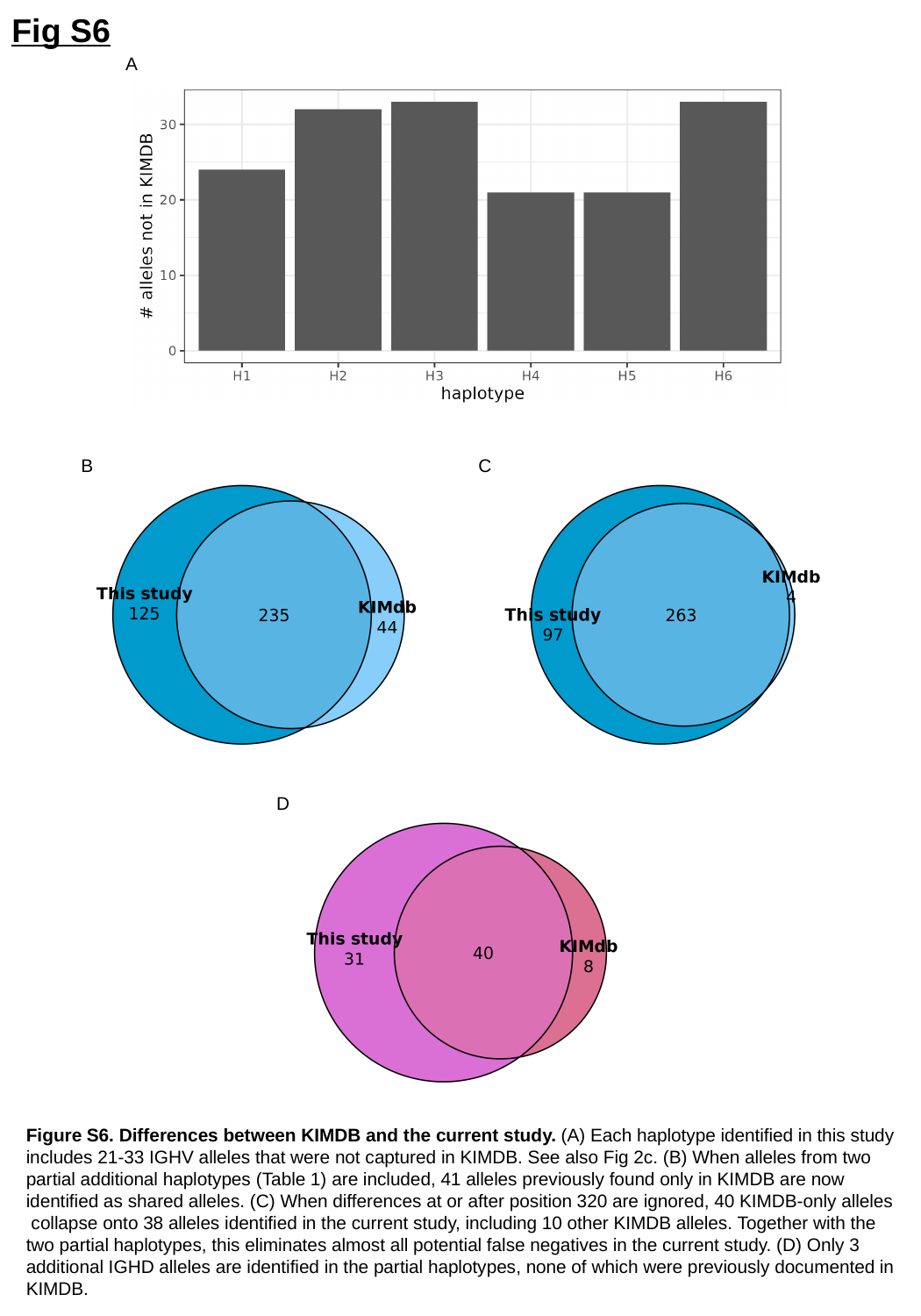

Fig S6
A
B
C
D
Figure S6. Differences between KIMDB and the current study. (A) Each haplotype identified in this study includes 21-33 IGHV alleles that were not captured in KIMDB. See also Fig 2c. (B) When alleles from two partial additional haplotypes (Table 1) are included, 41 alleles previously found only in KIMDB are now identified as shared alleles. (C) When differences at or after position 320 are ignored, 40 KIMDB-only alleles collapse onto 38 alleles identified in the current study, including 10 other KIMDB alleles. Together with the two partial haplotypes, this eliminates almost all potential false negatives in the current study. (D) Only 3 additional IGHD alleles are identified in the partial haplotypes, none of which were previously documented in KIMDB.
